# Dissolved organic matter from tropical peatlands impacts shelf sea light availability on coral reefs in the Singapore Strait, Southeast Asia

**DOI:** 10.1101/2021.03.30.437655

**Authors:** Patrick Martin, Nivedita Sanwlani, Tiffany Wan Qi Lee, Joel Meng Cheng Wong, Kristy Chang, Elizabeth Wing-See Wong, Soo Chin Liew

## Abstract

Shelf seas provide valuable ecosystem services, but their productivity and ecological functioning depend critically on sunlight transmitted through the water column. Anthropogenic reductions in underwater light availability are thus recognized as a serious threat to coastal habitats. The flux of strongly light-absorbing coloured dissolved organic matter (CDOM) from land to sea may have increased world-wide, but how this has altered the availability and spectral quality of light in shelf seas remains poorly known. Here, we present time-series data from the Sunda Shelf in Southeast Asia, where the monsoon-driven reversal in ocean currents supplies water enriched in CDOM from tropical peatlands for part of the year, resulting in 5–10-fold seasonal variation in light absorption by CDOM. We show that this terrigenous CDOM can dominate underwater light absorption at wavelengths up to 500 nm, and shift in the underwater irradiance spectrum towards longer wavelengths. The seasonal presence of terrigenous CDOM also causes the depth of 10% light penetration to shoal by 1–5 m, or 10–45%. We further estimate that on average 0.6 m, or 25%, of this terrigenous CDOM-mediated shoaling might be attributable to the enhanced loss of dissolved organic matter caused by peatland disturbance. We show that the seasonal change in the light environment is correlated with photo-acclimation by phytoplankton, and infer that terrigenous CDOM likely contributes to limiting the depth distribution of photosynthetic corals. Our results thus reveal an ecologically important but largely overlooked impact of human modifications to carbon fluxes that is likely becoming increasingly important in coastal seas.

## 1. Introduction

Shelf seas account for less than 10% of the global ocean area, but contribute more than 50% of the value of all marine ecosystem services (Costanza et al. 2014). These ecosystem services are largely contributed by benthic habitats that require sunlight for photosynthesis, such as coral reefs and seagrass beds. The attenuation of sunlight with depth is consequently a critical aspect of shelf sea water quality (Kirk 1988), and can strongly control the productivity and areal extent of light-dependent benthic ecosystems (Gattuso et al. 2006). Changes in light attenuation and water clarity are therefore highly significant stressors of shelf sea ecosystems (Dupont & Aksnes 2013, Filbee-Dexter & Wernberg 2018, Heery et al. 2018).

Underwater light attenuation varies chiefly as a result of absorption and backscattering of light by phytoplankton, suspended organic detritus particles, suspended inorganic sediment particles, and dissolved organic matter (DOM). Each of these optically active constituents can contribute significantly to light attenuation in aquatic ecosystems (IOCCG 2000). Moreover, because their absorption and backscattering spectra differ, the attenuation of an equal amount of sunlight by different constituents results in a different spectral distribution of light underwater. DOM absorbs most strongly at short wavelengths in the ultraviolet (UV) and blue range, and therefore plays an important role in protecting marine organisms from harmful UV radiation (Arrigo & Brown 1996, Banaszak & Lesser 2009, Häder et al. 2015). Yet by also absorbing blue light, DOM can shift the underwater irradiance to longer wavelengths that are less effectively absorbed by many photosynthetic organisms. Such spectral variation in the available light underwater can, for example, alter the phytoplankton community structure (Stomp et al. 2007, Gerea et al. 2017).

However, especially in the context of anthropogenic impacts on underwater light availability, the most widely recognized drivers of light attenuation in coastal waters are eutrophication (Dennison et al. 1993, Duarte 1995, Burkholder et al. 2007) and suspended sediment particles (Fabricius 2005, Storlazzi et al. 2015, Edmunds et al. 2018, Heery et al. 2018). In contrast, even though DOM can strongly control the optical properties of coastal waters (DeGrandpre et al. 1996, Kowalczuk et al. 2005, Foden et al. 2008, Kuwahara et al. 2010, Mascarenhas et al. 2017, Petus et al. 2018), the potential for DOM to drive ecologically significant changes in light attenuation, and the consequences for the spectral quality of underwater irradiance, are often neglected outside of the specialist optical oceanography literature.

The light absorbing substances in DOM, such as humic acids, are collectively known as coloured dissolved organic matter (CDOM). Although CDOM is also produced by bacteria and phytoplankton in the ocean (Coble 2007, Dainard & Guéguen 2013, Osburn et al. 2019), dissolved organic matter that originates from the partial decomposition of terrestrial vegetation is particularly strongly light-absorbent (Vantrepotte et al. 2015, Massicotte et al. 2017). The flux of this CDOM-rich terrigenous dissolved organic carbon (terrigenous DOC, or tDOC) from land to coastal oceans is a quantitatively significant part of the global carbon cycle and has increased by tens of percent in many parts of the world (Evans et al. 2005, Monteith et al. 2007, Moore et al. 2013). In Europe and North America, these trends may be largely driven by the recent reductions in atmospheric acid deposition, because high acid deposition can reduce organic matter solubility in soils (Skjelkvåle et al. 2005, Evans et al. 2006, Monteith et al. 2007). However, climate warming, increased water run-off, and land-use change are also driving increased tDOC fluxes both in high- and low-latitude regions (Hessen et al. 2010, Larsen et al. 2011, Weyhenmeyer et al. 2012, Moore et al. 2013, de Wit et al. 2016, Noacco et al. 2017, Wauthy et al. 2018). CDOM is an integral part of tDOC, and the concentrations of tDOC and of CDOM are therefore highly correlated in rivers and river plumes (Fichot & Benner 2011, Leech et al. 2016, Cook et al. 2017, Massicotte et al. 2017, Martin et al. 2018). Thus, increases in tDOC flux to coastal waters will necessarily entail increases in terrigenous CDOM flux. Moreover, tDOC and the associated terrigenous CDOM frequently exhibit relatively conservative mixing behaviour across salinity gradients in estuaries and shelf seas (Blough et al. 1993, Yamashita et al. 2011, Fichot & Benner 2012, Chen et al. 2015, Martin et al. 2018, Painter et al. 2018), such that shelf seas and adjacent oceanic regions can have high concentrations of tDOC and terrigenous CDOM (Blough et al. 1993, Kaiser et al. 2017, Medeiros et al. 2017, Carr et al. 2019, Zhou et al. 2019). Terrigenous CDOM thus has the potential to spread extensively across shelf seas, and increases in tDOC flux might therefore affect the light environment over large areas of coastal ocean.

That terrigenous, as opposed to marine, CDOM can significantly affect the underwater light environment in the sea has been recognised in regions where the CDOM pool is predominantly terrestrial (Blough et al. 1993, Kjeldstad et al. 2003, Kowalczuk et al. 2006, Hessen et al. 2010, Mizubayashi et al. 2013, Cherukuru et al. 2014). However, CDOM in coastal waters often consists of a mixture of marine and terrigenous CDOM in proportions that can vary strongly spatially and temporally. The most common way to distinguish marine from terrigenous CDOM is by measuring the slope of the CDOM absorption spectrum, originally over large wavelength ranges from UV to visible (Stedmon & Markager 2001, Kowalczuk et al. 2006, Astoreca et al. 2009). More recently, spectral slopes over narrow ranges of shorter wavelengths have become increasingly established for identifying terrigenous CDOM in terrestrially influenced marine waters, especially the spectral slope between 275–295 nm, S_275–295_ (Helms et al. 2008, Vantrepotte et al. 2015, Lu et al. 2016, Medeiros et al. 2017, Carr et al. 2019, Signorini et al. 2019). S_275–295_ has also been used successfully to quantify tDOC concentrations in shelf seas (Fichot & Benner 2012, Fichot et al. 2013, Fichot et al. 2014).

However, although the importance of terrigenous CDOM for the light environment of coastal waters has clearly been recognised, the relative contributions of terrigenous and marine CDOM to light attenuation in shelf seas have usually not been partitioned quantitatively. Consequently, although large-scale anthropogenic changes in land–ocean tDOC fluxes have the potential to alter the light environment of shelf seas by altering the terrigenous CDOM concentration, our knowledge of such impacts is still very limited. Based on long-term trends in salinity and correlations between salinity and CDOM in Norwegian fjords, Aksnes et al. (2009) inferred that increased terrigenous CDOM had resulted in “coastal browning”, which may have contributed to mesopelagic regime shifts from fish (visual predators) to jellyfish (tactile predators). Similarly, an increase in non-autotrophic particulate organic carbon in southern Norway was interpreted as indicating an increase in terrigenous CDOM, and this was hypothesised to have contributed to the collapse of coastal kelp forests (Frigstad et al. 2013). An increase in terrigenous CDOM was also invoked as a possible mechanism driving decreased water clarity in the North Sea (Dupont & Aksnes 2013), which may have caused a shift in the timing of the spring phytoplankton bloom by up to three weeks (Opdal et al. 2019). Ecosystem modelling has additionally demonstrated that increased terrigenous CDOM absorption can lead to a shallower distribution of phytoplankton and a shallower nutricline, resembling symptoms of eutrophication (Urtizberea et al. 2013). Moreover, recent time series of CDOM and comparison to historical chromaticity measurements suggest that terrigenous CDOM concentrations in the Gulf of Maine have increased as a result of greater river run-off (Balch et al. 2012, Balch et al. 2016), although the impacts for light attenuation could not be quantified in these studies. Consequently, it remains unclear to what extent coastal browning due to terrigenous CDOM has impacted shelf sea ecosystems and productivity.

This contrasts with our better understanding of the ecological impacts of freshwater “lake browning” caused by rising inputs of tDOC and terrigenous CDOM (Larsen et al. 2011, Graneli 2012, Wauthy et al. 2018). These impacts include reductions in primary productivity, shifts from benthic to pelagic primary productivity, thermocline shoaling, and possibly reductions in stocks of visually hunting fish (Ask et al. 2009, Thrane et al. 2014, Solomon et al. 2015, Vasconcelos et al. 2019). However, the freshwater lakes where lake browning has been reported contain far higher concentrations of terrigenous CDOM than do shelf seas, so much so that filtered lake water can have a noticeably yellow-brown colouration (Solomon et al. 2015). Whether CDOM-mediated browning can affect the ecological functioning of shelf seas to the same degree as lakes is still unclear.

Here, we use biogeochemical and optical time-series data from the Singapore Strait in the Sunda Shelf Sea in Southeast Asia to estimate seasonal changes in the proportion of marine and terrigenous CDOM, and then to quantify how terrigenous CDOM impacts underwater light availability. Tropical peatlands are the dominant source of tDOC and terrigenous CDOM in this part of Southeast Asia (Baum et al. 2007, Siegel et al. 2019). Previous research suggests that the extensive and recent anthropogenic disturbance and drainage of these peatlands (Miettinen et al. 2016) have increased their tDOC flux by about 50% (Moore et al. 2013, Yupi et al. 2016). Given that tDOC and CDOM show a strong relationship in Southeast Asian peatlands (Cook et al. 2017, Martin et al. 2018), we present a first-order estimate of the potential anthropogenic contribution to CDOM-mediated light attenuation.

## 2. Materials & Methods

### 2.1 Study area

The Singapore Strait is located in the central Sunda Shelf Sea, close to the peatlands on Sumatra (Fig. 1). The ocean currents on the Sunda Shelf reverse direction seasonally (van Maren & Gerritsen 2012, Mayer & Pohlmann 2014, Susanto et al. 2016, Wei et al. 2019): during the Northeast (NE) Monsoon (November–February), water flows westwards from the South China Sea through the Singapore Strait, and further northwards through the Malacca Strait to the Indian Ocean. During the Southwest (SW) Monsoon (May–September), the flow through the Malacca Strait stagnates or reverses direction, and water flows eastwards from the coast of Sumatra back through the Singapore Strait (Fig. 1). The annual mean flow is westwards from the South China Sea through the Singapore Strait and northward through the Malacca Strait to the Indian Ocean, with water residence times of 1–2 years for most parts of the shelf (Mayer et al. 2015, Mayer et al. 2018). This region of the Sunda Shelf (southern Malacca Strait, Singapore Strait, and Karimata Strait) experiences strong tidal currents that mix the water column all the way to the seafloor and prevent stratification (Mayer & Pohlmann 2014, Hamzah et al. 2020). This means that water from the open South China Sea reaches the Sumatran coast, receives large inputs of CDOM-rich tDOC from the peatlands, and then seasonally flows back into the Singapore Strait, while longer-term flowing towards the Indian Ocean *via* the Malacca Strait and the Java Sea. Seasonal variation in the Singapore Strait thus reflects primarily the regional spatial variability around the Singapore Strait. For the present analysis, we defined the seasons as follows: Intermonsoon 1: 01 March to 14 May; SW Monsoon: 15 May to 14 September; Intermonsoon 2: 15 September to 14 November; NE Monsoon: 15 November to end of February.

**Fig. 1.**
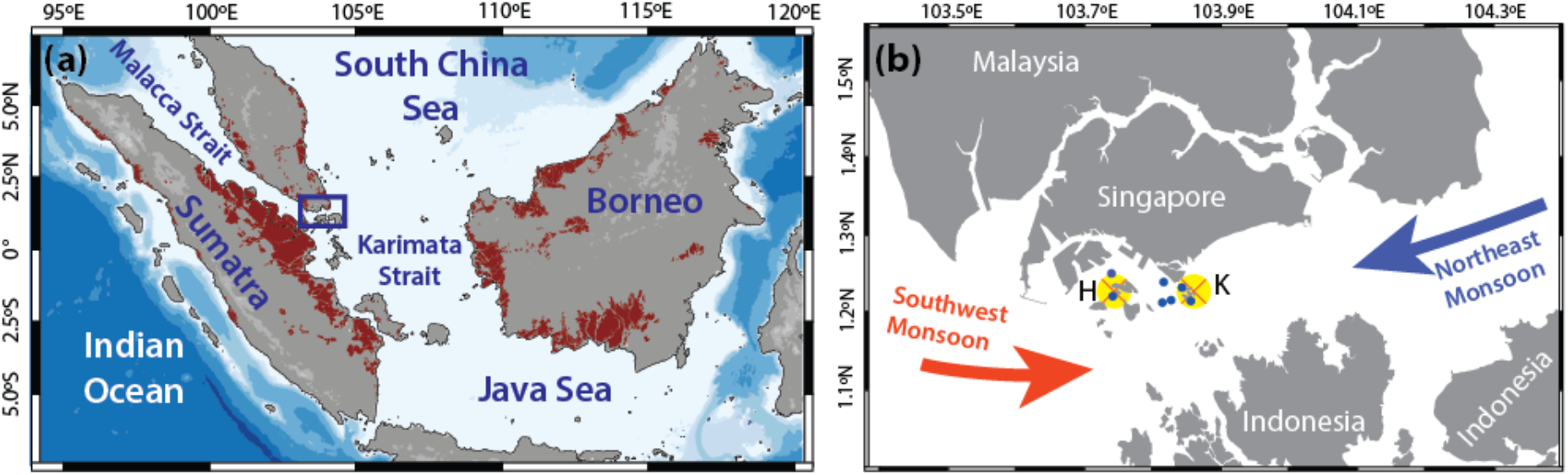
(a) Study region showing location of peatlands (brown shading) and ocean bathymetry; the purple box corresponds to the area shown in (b). (b) Map of the Singapore Strait, with the red and purple arrows showing the mean current direction during each monsoon season. The two main sampling sites are marked by the two crosses on yellow background; the eastern site (“K”) is the exposed site, Kusu, while the western site (“H”) is the sheltered site, Hantu. Other stations that were sampled occasionally are shown in small blue dots.

### 2.2 Field sampling and sensor measurements

The present analysis uses data collected as part of an on-going biogeochemical time-series programme. Measurements of salinity, DOC, and CDOM were collected at monthly to biweekly frequency between October 2017 and August 2020. Bio-optical parameters (particulate absorption and backscattering) were measured approximately monthly between December 2018 and August 2020. We collected samples at two sites (Fig. 1): Kusu Island (an exposed site experiencing higher wave energy) and Hantu Island (a sheltered site with lower wave energy). Both islands have artificial breakwaters, and narrow coral reefs that extend about 20 m horizontally from the breakwaters. Additional sites in between Kusu and Hantu were sampled occasionally to constrain spatial variability (Fig. 1). Conductivity-temperature-depth profiles were measured with a Valeport FastCTD at 16 Hz to 12–15 m depending on current and bottom depth; stratification was not observed. Water was collected adjacent to the reefs at 5 m depth using a Niskin bottle. Samples for CDOM and DOC analysis were filtered on board through 0.2 μm Supor polyethersulfone filters (all tubing and filter housings were cleaned with 1 M HCl, then assembled with the filters and pre-rinsed with 200 mL of ultrapure water (18.2 MΩ cm^−1^) and with sample water;) and stored in 40-mL amber borosilicate EPA vials (pre-baked at 450°C for 4 h) with Teflon-coated septa. Water for chlorophyll-*a* and particulate absorption was stored in acid-washed HDPE bottles in the dark and filtered (25-mm Whatman GF/F) 3–6 hours later in the laboratory.

Backscattering was measured at 412, 440, 488, 510, 532, 595, 650, 676, and 715 nm using a Wetlabs BB9 lowered to 1 m depth; 60 consecutive measurements were taken at 1 Hz and averaged. The data were processed according to manufacturer instructions: the raw measurement was converted to the total volume scattering coefficient by subtracting the dark offset (measured before each field trip) and multiplying by a calibration scaling factor, and corrected for non-water absorption as measured by a TriOS OSCAR instrument. Volume scattering from pure seawater was subtracted and the particulate scattering converted to particulate backscattering coefficients following Boss and Pegau (2001), which were then fit with a power law at 1-nm resolution over the wavelength range of photosynthetically active radiation (PAR, 400–700 nm).

The present analysis focuses on the period of December 2018 to August 2020, during which we acquired 77 measurements of CDOM, DOC, and salinity, 60 measurements of particulate absorption, and 36 measurements of backscattering.

### 2.3 CDOM and particulate absorption measurements

CDOM samples were stored at +4°C back on land and analysed within 24 h of collection. Samples were brought to room temperature and absorbance measured from 250–800 nm at 1-nm resolution in 10-cm pathlength quartz cuvettes on a Thermo Evolution300 dual-beam spectrophotometer against ultrapure water as a reference. Data were baseline-corrected according to Green and Blough (1994), smoothed using a loess function, and converted to Napierian absorption coefficients, using the R package hyperSpec (Beleites & Sergo 2018). Here, we express the concentration of CDOM as the CDOM absorption coefficient at 440 nm, *a*_CDOM_(440), with units of m^−1^. We also calculated the CDOM spectral slope from 275–295 nm, S_275-295_, and the spectral slope ratio (S_R_, the ratio of the 275–295 nm slope to the 350–400 nm slope) following Helms et al. (2008). Both S_275–295_ and S_R_ have been shown to correlate with DOM molecular weight (Helms et al. 2008) and are widely used as markers of tDOC in coastal seas (Fichot & Benner 2012, Fichot et al. 2013, Vantrepotte et al. 2015, Lu et al. 2016, Medeiros et al. 2017, Painter et al. 2018, Carr et al. 2019). To test whether seasonal variation in CDOM could be explained by conservative mixing between terrigenous CDOM (with low S_275–295_ and S_R_) and marine CDOM (with high S_275–295_ and S_R_), we calculated a theoretical mixing model between the CDOM spectra measured on 15 March 2019 (with high S_275–295_ and low *a*_CDOM_(440)) and 16 July 2020 (lowest S_275–295_ and highest *a*_CDOM_(440) in 2019 and 2020) as follows:

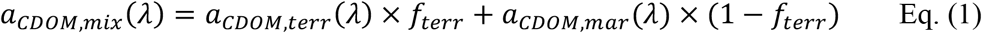

where *a*_CDOM,mix_(*λ*) is the predicted CDOM spectrum for conservative mixing, *a*_CDOM,terr_ and *a*_CDOM,mar_ are the measured CDOM spectra at Hantu Island on 16 July 2020 and 15 March 2019, respectively, and *f*_terr_ is the fractional contribution of the 16 July 2020 spectrum (which we varied between 0 and 1 in increments of 0.0125 to simulate conservative mixing). For each predicted *a*_CDOM,mix_ spectrum, S_275-295_ and S_R_ were recalculated as for the original data. The theoretical mixing curves were plotted together with the measured data on scatter plots of S_275–295_ and S_R_ against *a*_CDOM_(440), following Stedmon and Markager (2001).

Samples for particulate absorption (500–1000 mL) were vacuum-filtered onto 25-mm diameter Whatman GF/F filters and stored in liquid nitrogen in tissue embedding cassettes (Kartell Labware) wrapped in aluminium foil. Samples were thawed to room temperature, moistened by briefly placing them on a sponge soaked in filtered seawater, and absorbance measured from 300–800 nm with filters held inside an integrating sphere using a centre-mount sample holder on a PerkinElmer Lambda 950 spectrophotometer, as recommended by Stramski et al. (2015). Multiple blank filters were measured throughout each batch of analysis. Filters were then depigmented (as assessed by the complete disappearance of the chlorophyll-*a* absorption peak at 668 nm) with 5 ml of 0.1% sodium hypochlorite in ultrapure water with 60 g l^−1^ sodium sulphate for 15 min (Ferrari & Tassan 1999), rinsed with 5 ml ultrapure water, and remeasured. All blank and sample absorbance spectra were first corrected for baseline drift by subtracting the mean absorbance from 801–851 nm from the rest of the spectrum, and then blank-corrected by subtracting the mean baseline drift-corrected blank spectrum from all sample absorbance spectra. These corrected sample absorbances were then corrected for pathlength amplification according to Stramski et al. (2015):

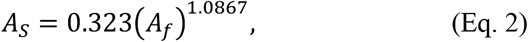

where *A*_*s*_ is the pathlength-corrected sample absorbance, and *A*_*f*_ is the blank- and baseline-corrected absorbance of each filter. Corrected absorbances were then converted to Napierian absorption coefficients by accounting for the area of sample on each filter (the filtered area of each filter had a radius of 11.5 mm) and the sample volume filtered. Phytoplankton absorption (*a*_phyto_) was calculated by subtracting the depigmented absorption spectrum (i.e., the non-algal particulate absorption, *a*_NAP_) from the total particulate absorption spectrum. From the *a*_phyto_ spectra, we further calculated the phytoplankton absorption spectral slope following Eisner et al. (2003):

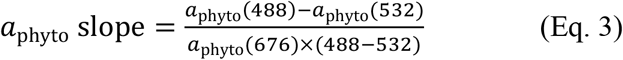

where numbers in parentheses indicate the wavelengths. We also calculated the ratio of phytoplankton absorption at 490 to 510 nm, *a*_phyto_(490):*a*_phyto_(510), following Hickman et al. (2009). Both of these are measures of the ratio of photoprotective to photosynthetic carotenoid pigments in the phytoplankton community, which is indicative of photo-acclimation.

Seasonal average spectral absorption budgets were calculated as the fractional contribution of each absorbing constituent (CDOM, non-algal particles, phytoplankton, and water) to the total absorption at each wavelength, and then averaging these data seasonally.

### 2.4 DOC analysis and specific UV absorbance calculation

DOC samples (30 ml) were acidified with 100 μl of 50% H_2_SO_4_ in the field, stored at +4°C, and analysed within 2–3 months of collection on a Shimadzu TOC-L analyser with the Shimadzu high-salt combustion kit and calibrated using potassium hydrogen phthalate, as in our previous work (Martin et al. 2018). Certified reference material from the University of Miami (deep-sea water, 42–45 μmol l^−1^ DOC) was analysed alongside every batch of measurements, and returned a long-term mean ± standard deviation of 48.0 ± 3.9 μmol l^−1^ throughout our time series. DOC samples were collected and analysed in triplicate starting in December 2018. We used these data to calculate the specific UV absorbance at 254 nm (SUVA_254_):

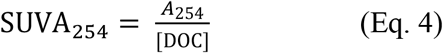

where *A*_254_ is the CDOM absorbance at 254 nm per metre, and the DOC concentration is in mg C l^−1^ (note that the absorbance is obtained by dividing the Napierian absorption coefficient by 2.303). SUVA_254_ consequently has units of l mg^−1^ m^−1^, and is a measure of DOM aromaticity (Traina et al. 1990, Weishaar et al. 2003). Similar to S_275–295_ and S_R_, SUVA_254_ is useful as a tracer of terrigenous DOM in aquatic ecosystems (Cao et al. 2018, Anderson et al. 2019, Carr et al. 2019).

### 2.5 Chlorophyll-*a*

Samples for chlorophyll-*a* (200–1000 ml) were filtered onto 25 mm diameter Whatman GF/F filters, wrapped in aluminium foil, flash-frozen in liquid nitrogen, and stored at −80°C until analysis within 3 months. Filters were then extracted in 90% acetone at 4°C in the dark overnight, briefly centrifuged to remove particles, and fluorescence measured on a Horiba Fluoromax4 at excitation 436 nm and emission 680 nm with slit widths of 5 nm (Welschmeyer 1994). Fluorescence was acquired as the fluorescence signal normalised to the lamp reference measurement to account for variation in lamp intensity (using the Fluoromax4 S1c/R1c acquisition mode), and calibration was performed with a spinach chlorophyll-*a* standard (Sigma-Aldrich, C5753-1MG).

### 2.6 Calculating light attenuation spectra

Underwater light attenuation can be described by the diffuse attenuation coefficient of downwelling irradiance, *K*_*d*_, which varies spectrally:

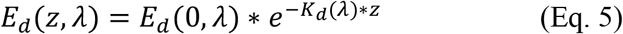

where *E*_*d*_(*z,λ*) is the downwelling irradiance at depth *z* and wavelength *λ*, *E*_*d*_(0*,λ*) is the downwelling irradiance at wavelength *λ* at the surface, and *K*_*d*_(*λ*) is the diffuse attenuation coefficient at wavelength *λ*. We used our spectral measurements of absorption by CDOM and particles, and backscattering by particles, to calculate spectra of *K*_*d*_ over the wavelength range 400–700 nm according to Lee et al. (2005):

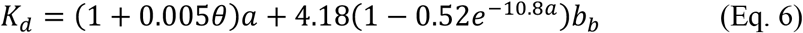

where *θ* is the solar zenith angle, *a* is total absorption, and *b*_*b*_ is total backscattering. Absorption and backscattering spectra of pure seawater were taken from Pope and Fry (1997) and Smith and Baker (1981), respectively. We used the solar zenith angle at solar noon on each date (i.e., the time of day when the sun is at its highest point), such that the result reflects the maximum light penetration for each date. Solar zenith angles and solar noon times were calculated using the R packages GeoLight (Lisovski & Hahn 2012) and suncalc. This calculation was originally developed to estimate *K*_*d*_ between the surface and the depth to which 10% of surface PAR penetrates (*Z*_10%_) and was therefore denoted *K*_*d*_(*E_10%_*) by Lee et al. (2005); we refer to this as *K*_*d*_ here for simplicity, since *K*_*d*_ at individual wavelengths does not vary strongly with depth unless the absorption and backscattering spectra vary with depth. A total of 32 concomitant measurements of absorption and backscattering were available for this calculation, taken on 17 separate dates. We verified that this calculation yielded accurate estimates of *K*_*d*_ by comparing calculated *K*_*d*_ spectra to *K*_*d*_ spectra measured using a TriOS RAMSES radiometer at 19 of these 32 stations; the radiometer measurement methods and the results of this comparison are shown in the Supplementary Information.

### 2.7 Calculating underwater irradiance spectra and depth of PAR penetration

To examine how seasonal variation in absorption and backscattering affect both the spectral quality of irradiance underwater and the depth to which PAR penetrates, we used the *K*_*d*_ spectra together with modelled mid-day solar irradiance for each date to calculate depth profiles of underwater irradiance, the vertical attenuation coefficient of downwelling PAR, *K*_*d*_(PAR), and *Z*_10%_. Our objective with this analysis was not to derive the actual underwater irradiance on each date, which depends especially on cloud cover, but rather to determine the potential effects of the observed variation in absorption and backscattering on the underwater light environment. We therefore modelled the downwelling irradiance spectrum just below the water surface (*E*_*d*_0^−^) for solar noon on each date using the Hydrolight model, assuming identical cloud cover and wind speed for each day (20% and 2 m s^−1^, respectively), and used these modelled spectra as inputs for our calculations. This means that seasonal changes in solar zenith angle (and their resulting effects on irradiance) are accounted for, but that our results are otherwise representative of conditions experienced around mid-day on relatively cloud-free days. Variation in cloud cover chiefly alters the total irradiance, but does not affect the shape of the irradiance spectrum very strongly. Our purpose with these calculations was not to estimate exact light doses, but rather to examine how the depth penetration and spectral distribution of underwater light vary over time as a result of our measured changes in absorption and backscattering, for which modelled irradiances are sufficient. We first calculated the average underwater irradiance spectrum experienced by phytoplankton in a fully mixed water column, *E*_*d*_(*Z*_mean_) (Ferrero et al. 2006, Gerea et al. 2017):

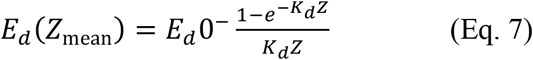

where *Z* is the depth of the water column. We selected 30 m, which is representative of much of the Singapore Strait surrounding our sampling sites (Chan et al. 2006).

Next, to examine how the spectral light quality experienced by benthic organisms is affected, we calculated the underwater irradiance spectrum at fixed depths within the upper 10 m for each date by attenuating the Hydrolight-modelled noon-time *E*_*d*_0^−^ spectra with the calculated *K*_*d*_ spectra according to Eq. (5).

Finally, to examine how the overall depth of light penetration varies, we calculated *K*_*d*_(PAR) and *Z*_10%_. To do this, we first attenuated the modelled *E*_*d*_0^−^ spectra with the calculated *K*_*d*_ spectra (Eq. 5) at 0.1 m intervals from the surface down to a depth of 20 m to yield calculated depth profiles of downwelling irradiance (*E*_*d*_). The calculated *E*_*d*_ spectrum at each depth was then converted from W m^−2^ nm^−1^ to the downward flux of photons, *E*_*q*_, at each wavelength according to:

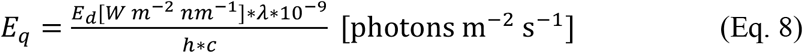

where *h* is Planck’s constant and *c* is the speed of light in m s^−1^. *E*_*q*_ was converted to μmol photons m^−2^ s^−1^ and then summed across the wavelength range of 400–700 nm to yield a quantum flux of PAR at each depth. *K*_*d*_(PAR) was then calculated as the slope of a linear regression of the natural log of quantum PAR flux *versus* depth, and *Z*_10%_ was calculated as 2.303/*K*_*d*_(PAR). Unlike *K*_*d*_ at individual wavelengths, *K*_*d*_(PAR) changes significantly with depth because of the large spectral variation in *K*_*d*_(λ) (Lee 2009, Lee et al. 2018).

Consequently, the value of *K*_*d*_(PAR) calculated by regressing PAR against depth varies depending on the depth to which the regression is performed. Since our objective with this calculation was to quantify *Z*_10%_, the regression should ideally be performed down to *Z*_10%_ rather than to a fixed, arbitrary depth, so we sought to first estimate the approximate depth of *Z*_10%_ to determine the appropriate depth to which to perform the regression. Using our 19 measured radiometer profiles (described in the Supplementary Information), we found that *Z*_10%_ was closely related to *K*_*d*_ at 520 nm:

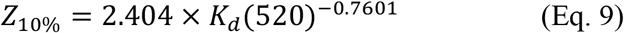

where *K*_*d*_(520) is *K*_*d*_ at 520 nm (Fig. S1). We used this initial estimate of *Z*_10%_ for each station as the depth over which we calculated *K*_*d*_(PAR) using a PAR *versus* depth regression as explained above. The final value of *Z*_10%_ for each station was then calculated from *K*_*d*_(PAR) as described above.

### 2.8 Impact of terrigenous CDOM on *Z*_10%_

Our time-series of S_275–295_, S_R_, and SUVA_254_ indicated that the variation in CDOM absorption is predominantly the result of conservative mixing between terrigenous CDOM and marine CDOM, as shown in Section 3.1 below. Based on these data, the CDOM during the March–April intermonsoon period was predominantly marine, while the CDOM during other periods consisted of a mixture of this background level of marine CDOM and a varying amount of terrigenous CDOM. We therefore quantified the amount of terrigenous CDOM in each sample by subtracting the intermonsoon CDOM spectrum measured on 15 March 2019 (which we also used as one endmember in our conservative mixing model; see Section 2.3) from the measured CDOM spectrum in each sample.

To quantify the impact of this terrigenous CDOM on the depth of PAR penetration, we recalculated our *K*_*d*_ spectra (Eq. 6) using the 15 March 2019 CDOM spectrum in place of the CDOM spectrum measured for each station. We then recalculated *K*_*d*_(PAR) and *Z*_10%_, as well as *E*_*d*_(*Z*_mean_), as described in Section 2.7. This yielded estimates of what *K*_*d*_(PAR), *Z*_10%_, and *E*_*d*_(*Z*_mean_) would have been at each station in the absence of terrigenous CDOM.

To quantify the potential anthropogenic contribution to CDOM-mediated light attenuation, we recalculated *K*_*d*_ again, this time with the terrigenous CDOM absorption reduced by 35% of the observed value. This is based on estimates from Borneo and Sumatra that land-use change has increased the flux of DOC from Southeast Asian peatlands by 54% (Moore et al. 2013, Yupi et al. 2016), and the fact that nearly all peatlands in the region have experienced disturbance (Miettinen et al. 2016). Because DOC and CDOM are very closely correlated in peatland-draining blackwater rivers and downstream coastal waters in Southeast Asia (Cook et al. 2017, Martin et al. 2018), these estimates imply that 35% of the observed terrigenous CDOM in peatland-influenced coastal waters is anthropogenic (if the modern, post-disturbance DOC flux is 1.54-fold greater than the pre-disturbance DOC flux, then the anthropogenic fraction of the modern DOC flux is 0.54/1.54 = 0.35). We only estimated this anthropogenic contribution for the SW Monsoon period, as this is the season when the Singapore Strait receives terrestrial inputs from the large peatland areas on Sumatra.

## 3. Results

### 3.1 Bio-optical time-series data

The concentration of CDOM, *a*_CDOM_(440), ranged from lowest values of 0.039–0.045 m^−1^ during the intermonsoon seasons to peak values of 0.27–0.45 m^−1^ during the May–September SW Monsoon (Fig. 2a). Smaller increases were also seen during the November–February NE Monsoon, with peak *a*_CDOM_(440) of 0.10–0.17 m^−1^. During the SW Monsoon, the CDOM spectral slope between 275–295 nm (S_275–295_) decreased from around 0.030 to ≤0.018 (Fig. 2b), and the spectral slope ratio (S_R_) decreased from values around 2.0 to <1.25 (Fig. 2c), while the SUVA_254_ increased from values <1.0 to mostly between 2.0–3.0 (Fig. 2d). Smaller decreases in S_275–295_ and S_R_, and increases in SUVA_254_, were also seen during the NE Monsoon. Seawater salinity decreased from values of 32–33 during the intermonsoon periods to 31–32 during the NE Monsoon and even lower to 29–31 during the SW Monsoon (Fig. 2e). Additional data collected in 2018 show that the seasonal increases in CDOM and SUVA_254_, and decreases in salinity, S_275-295_, and S_R_ were similar to 2020, confirming that large and sustained inputs of CDOM are typical during the SW Monsoon (Fig. S2; particulate optical properties were not measured until 2019). These seasonal differences in *a*_CDOM_(440), S_275–295_, S_R_, SUVA_254_, and salinity were all statistically significant (Kruskal-Wallis test, all *χ*^2^ >77, d.f. = 3, all *p* <0.001)

**Fig. 2.**
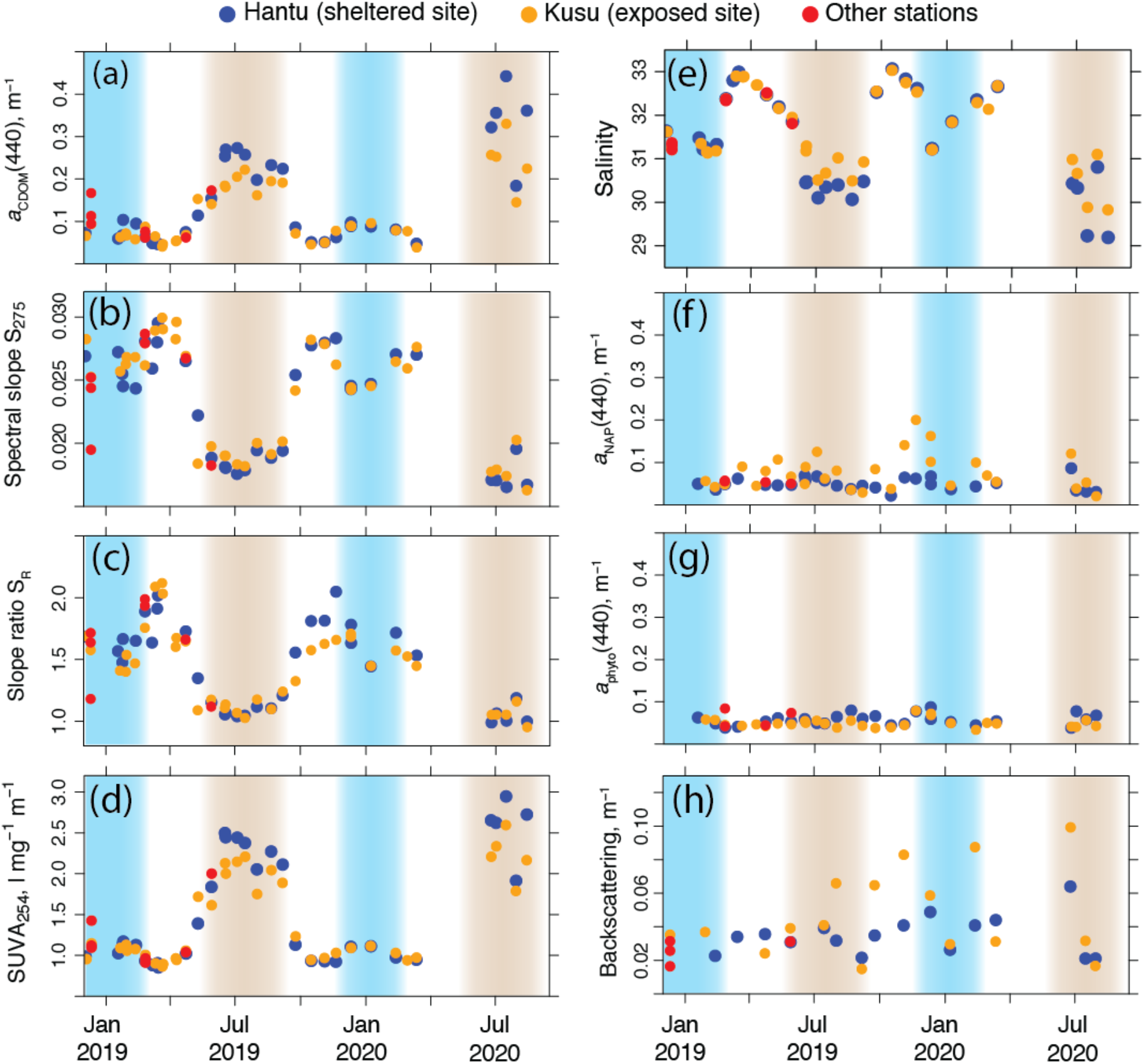
Biogeochemical and optical time-series data. (a) Absorption coefficient of CDOM at 440 nm, (b) CDOM spectral slope between 275–295 nm, (c) CDOM spectral slope ratio, (d) specific UV absorbance at 254 nm (SUVA_254_), and (e) seawater salinity all showed seasonal variation consistent with a large input of terrigenous CDOM during the Southwest Monsoon (brown shading), and to a lesser extent during the Northeast Monsoon (blue shading). In contrast, time series of absorption by (f) non-algal particles and (g) phytoplankton (both at 440 nm), and (h) backscattering by particles (at 440 nm) showed no clear seasonality, but absorption by non-algal particles and particulate backscattering were quite variable and typically higher at the more exposed site.

Absorption by phytoplankton, non-algal particles, and particulate backscattering showed no significant differences between seasons (Kruskal-Wallis test, all *χ*^2^ <6.2, d.f. = 3, all *p* >0.10), although the more exposed site typically had higher non-algal particulate absorption and particulate backscattering (Figs. 2f–h). Note that particulate backscattering at 440 nm was highly correlated with particulate backscattering at each of the other eight wavelengths (Fig. S3; all Pearson’s correlation coefficients >0.97). Consistent with the low phytoplankton absorption, chlorophyll-*a* concentrations were relatively low (mean ± standard deviation of 1.0 ± 0.5 μg l^−1^; Fig. S4) and did not show significant differences between seasons (Kruskal-Wallis test, *χ*^2^ = 2.9, d.f. = 3, *p* = 0.40).

We found that our measured S_275–295_ and S_R_ values showed tightly constrained relationships with *a*_CDOM_(440) across all seasons (Fig. 3a,b), which closely followed the theoretical mixing model (Eq. 1; grey lines in Fig. 3a,b) between the CDOM spectra measured on 15 March 2019 (intermonsoon) and on 16 July 2020 (SW Monsoon). Note that CDOM spectral slope parameters show non-linear changes during conservative mixing (Stedmon & Markager 2003). Moreover, there was a strong, positive correlation between *a*_CDOM_(440) and SUVA_254_ (Fig. 3c; Spearman’s rank correlation, *ρ* = 0.906, *p* < 0.001, *n* = 129).

**Fig. 3.**
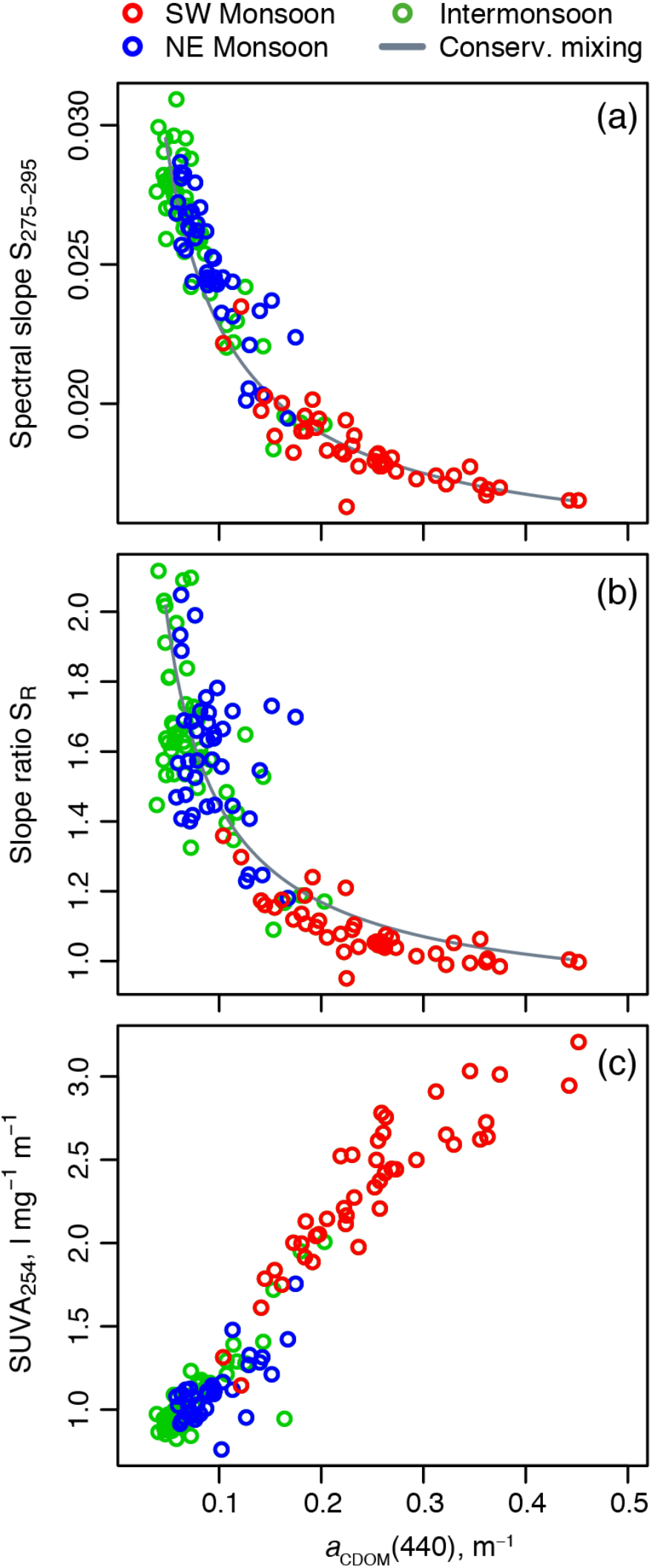
Scatterplots showing strong relationships between (a) S_275–295_, (b) S_R_, and (c) SUVA_254_ and CDOM absorption at 440 nm. Grey lines in (a,b) show predicted variation from the conservative mixing between the marine CDOM spectrum measured on 15 March 2019 and the primarily terrigenous CDOM spectrum measured on 16 July 2020. Data from the SW Monsoon consistently show high CDOM absorption associated with low S_275–295_, low S_R_, and high SUVA_254_, indicative of a primarily terrigenous CDOM pool during this season.

### 3.2 Seasonal changes in light absorption budgets, underwater irradiance, and phytoplankton absorption spectra

The large seasonal changes in CDOM absorption altered the average spectral light absorption budget between seasons (Fig. 4). Absorption in the UV range (300–400 nm) was dominated by CDOM in all seasons, but to a greater extent in the SW Monsoon. Between 400–500 nm, CDOM was progressively less dominant, and especially in the intermonsoon seasons the absorption by CDOM at 500 nm was only around 20% of the total absorption (Fig. 4a,c). In the SW Monsoon, however, CDOM contributed ≥50% of the total absorption up to 500 nm, and still contributed 50% of the non-water absorption up to 600 nm (Fig. 4b). During the NE Monsoon, the absorption budget was less CDOM-dominated than in the SW Monsoon, but more than during the intermonsoon seasons (Fig. 4e). In all seasons, absorption by non-algal particles was greater than phytoplankton absorption from 300 nm to roughly 440 nm, then up to 500–550 nm phytoplankton and non-algal particles contributed roughly equally, beyond which phytoplankton increasingly dominated the particulate absorption. In all seasons, water contributed >50% of absorption upwards of 550–570 nm (Fig. 4).

**Fig. 4.**
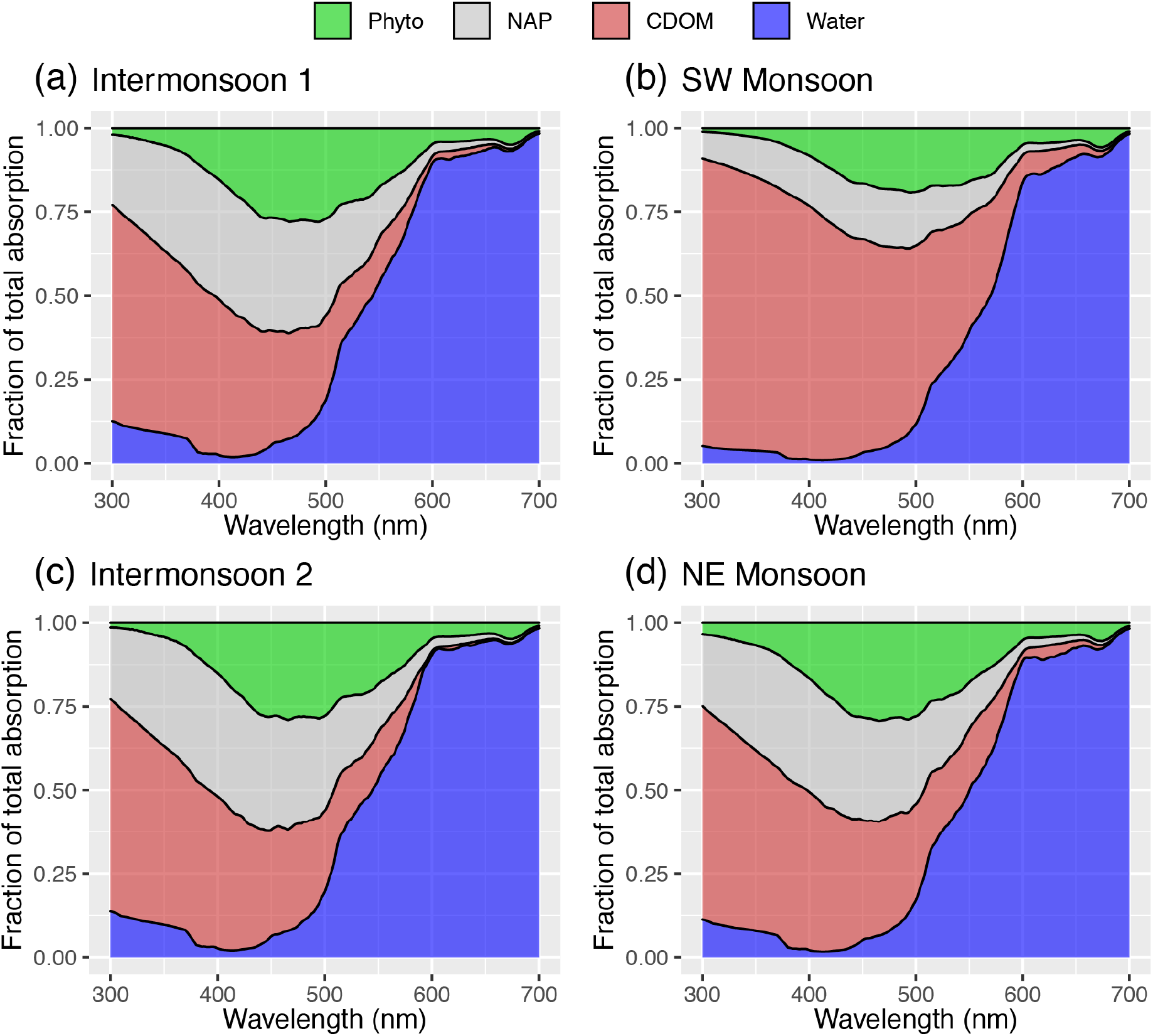
Light absorption budgets across the ultraviolet and visible wavelengths. Data are seasonal means across all stations. Phyto = absorption by phytoplankton; NAP = absorption by non-algal particles; CDOM = absorption by coloured dissolved organic matter; Water = absorption by seawater.

Using the modelled surface irradiance for solar noon on each sampling date, we found that the underwater irradiance at fixed depths between 1 and 10 m was shifted to longer wavelengths in the SW Monsoon: the wavelength of peak irradiance ranged from 531–539 nm during the intermonsoon and NE Monsoon seasons, but was shifted to between 547–566 nm in the SW Monsoon (Fig. 5). Similarly, the ratio of blue to green irradiance (calculated as *E*_d_(440) to *E*_d_(550)) at each depth was significantly lower during the SW Monsoon than other seasons (Kruskal-Wallis test, all *χ*^2^ ≥13.5, d.f. = 3, all *p* <0.005), indicating a seasonal decrease in the availability of blue light relative to longer wavelengths.

**Fig. 5.**
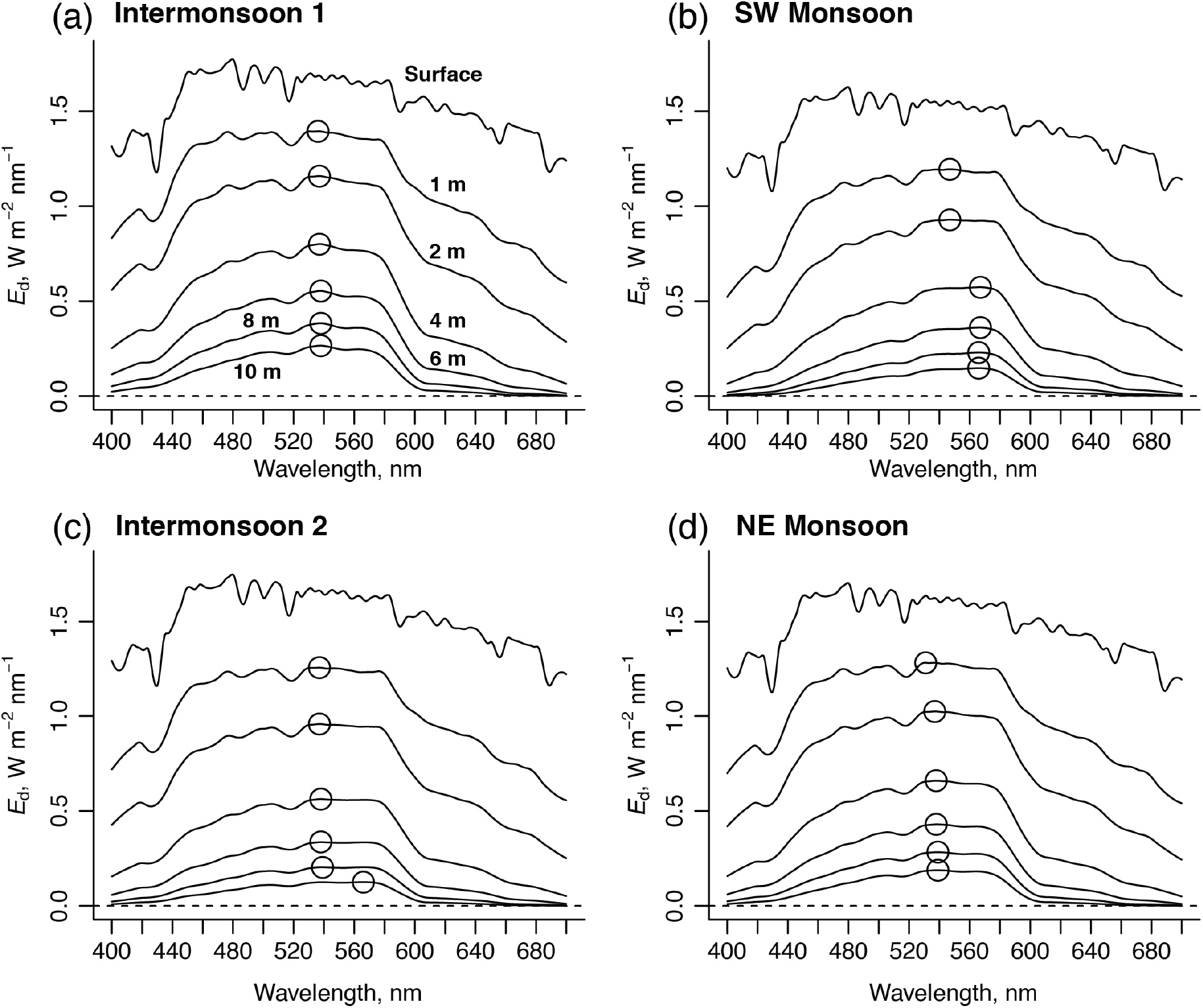
Seasonal average downwelling irradiance (*E*_d_) spectra calculated for solar noon at fixed depths for (a) the March–April Intermonsoon 1; (b) the Southwest Monsoon; (c) the October–November Intermonsoon 2; and (d) the Northeast Monsoon. Circles indicate the wavelength of maximum irradiance for each spectrum. The depths for spectra in panels c–d are as in panel a.

This spectral shift in the underwater irradiance was also evident in the average irradiance at solar noon experienced by phytoplankton under turbulent mixing (Eq. 7), which peaked at 567 nm during the SW Monsoon, but at 537–538 nm during the other seasons (Fig. 6). The ratio of blue to green irradiance was also significantly lower during the SW Monsoon (mean ratio of 0.46) compared to the other seasons (mean ratios of 0.54–0.62) for these averaged irradiances (Kruskal-Wallis test, *χ*^2^ = 25.9, d.f. = 3, *p* <0.001).

**Fig. 6.**
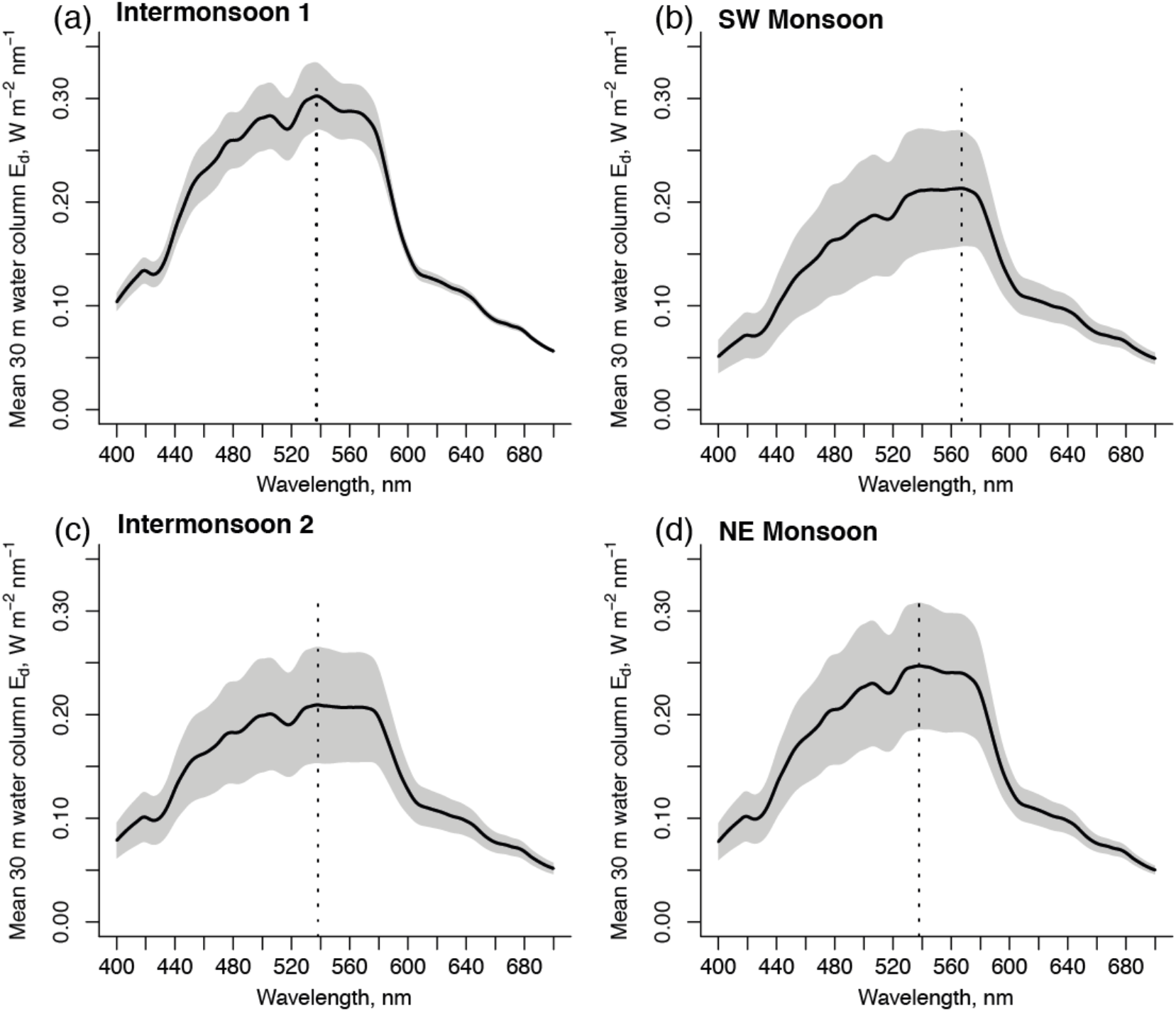
Seasonal averages of depth-averaged downwelling irradiance at solar noon from 0–30 m, assuming turbulent mixing of the water column, for (a) March–April Intermonsoon 1, (b) Southwest Monsoon, (c), October–November Intermonsoon 2, and (d) Northeast Monsoon. Solid black line indicates the seasonal average irradiance spectrum, grey shading indicates ± 1 standard deviation. Vertical dotted lines indicate the wavelength of maximum irradiance.

The phytoplankton absorption spectra revealed a statistically significant decrease in the ratio of *a*_phyto_(490) to *a*_phyto_(510) and a significant increase in the *a*_phyto_ spectral slope during the SW Monsoon (Fig. 7; Kruskal-Wallis test; both *χ*^2^ ≥12.4, d.f. = 3, both *p* ≤0.006). Both the ratio of *a*_phyto_(490) to *a*_phyto_(510) and the *a*_phyto_ spectral slope were also significantly correlated with *a*_CDOM_(440) (Spearman’s rank correlation; *ρ* = −0.45 and 0.39; both *p* <0.003) and with S_275–295_ (Spearman’s rank correlation; *ρ* = 0.50 and −0.43; both *p* <0.001).

**Fig. 7.**
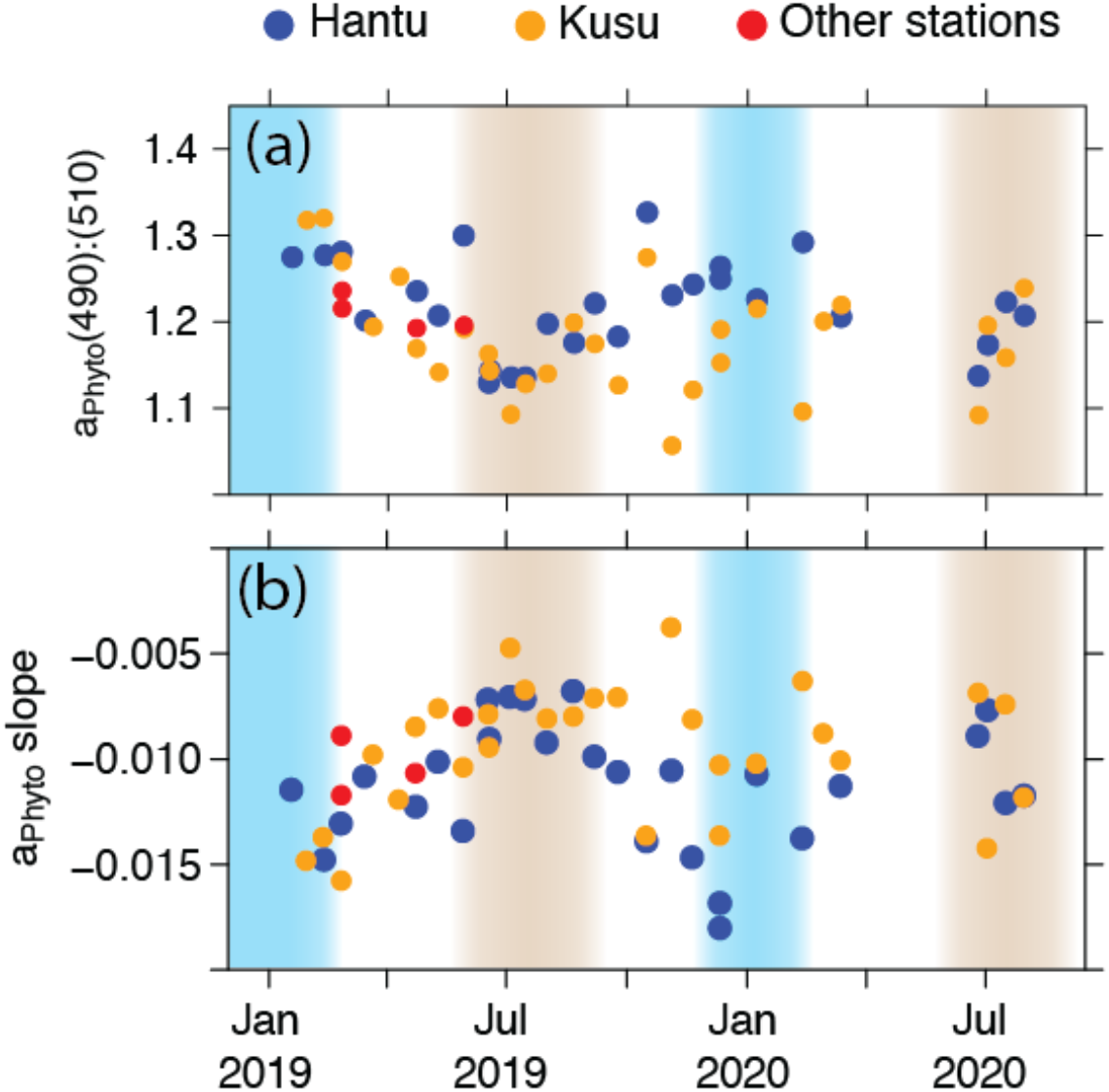
Time series of (a) the ratio of phytoplankton absorption at 490 nm to phytoplankton absorption at 510 nm, and (b) the phytoplankton absorption spectral slope (Eisner et al. 2003) both indicate that phytoplankton during the Southwest Monsoon have a lower proportion of non-photosynthetic to photosynthetic carotenoid pigments than during the other seasons (seasonal shading is as in Figs. 2 and 8).

### 3.3 *Z*_10%_ and impacts of terrigenous CDOM

The depth of 10% PAR penetration, *Z*_10%_, ranged between 3.7–9.8 m, with an overall average of 7.1 m, and was typically deeper at the more sheltered site (Fig. 7a). Across both sites, *Z*_10%_ was on average deepest during Intermonsoon 1 (8.6 m) and shallowest during the SW Monsoon (6.6 m); this seasonal difference was statistically significant (Kruskal-Wallis test, *χ*^2^ = 8.4, d.f. = 3, *p* = 0.038). To quantify the impact of terrigenous CDOM on the euphotic zone depth, we recalculated *K*_*d*_(PAR) and *Z*_10%_ by using only the background marine CDOM spectrum measured on 15 March 2019 in place of the observed total CDOM absorption (see Section 2.8). We found that relative to the actually observed value of *Z*_10%_, the value of *Z*_10%_ without terrigenous CDOM was deeper by 0.7–4.9 m (on average, 2.4 m). This corresponded to a shoaling of *Z*_10%_ by 13–45% due to terrigenous CDOM (Fig. 8a,b). Even during the NE Monsoon, when the terrigenous CDOM concentration was lower than during the SW Monsoon, the euphotic zone was shoaled by up to 1.9 m, or 17%. Across all seasons and sites, the percentage shoaling of *Z*_10%_ was strongly related to the terrigenous CDOM concentration (Fig. 8c), which shows that terrigenous CDOM had a large impact on the depth of underwater light penetration despite variation in the concentrations of suspended sediments and phytoplankton between sites and dates. We also repeated our calculation of the depth-averaged irradiance in a turbulent water column (Eq. 7), but using the *K*_*d*_ spectra calculated without terrigenous CDOM (see Section 2.8). We found that without terrigenous CDOM, the wavelength of peak irradiance was on average nearly identical between seasons (531–534 nm), and that the blue-to-green ratio *E_d_*(440): *E_d_*(550) no longer showed significant seasonal differences (average of 0.59–0.62 for each season; Kruskal-Wallis test, *χ*^2^ = 1.9, d.f. = 3, *p* = 0.599).

**Fig. 8.**
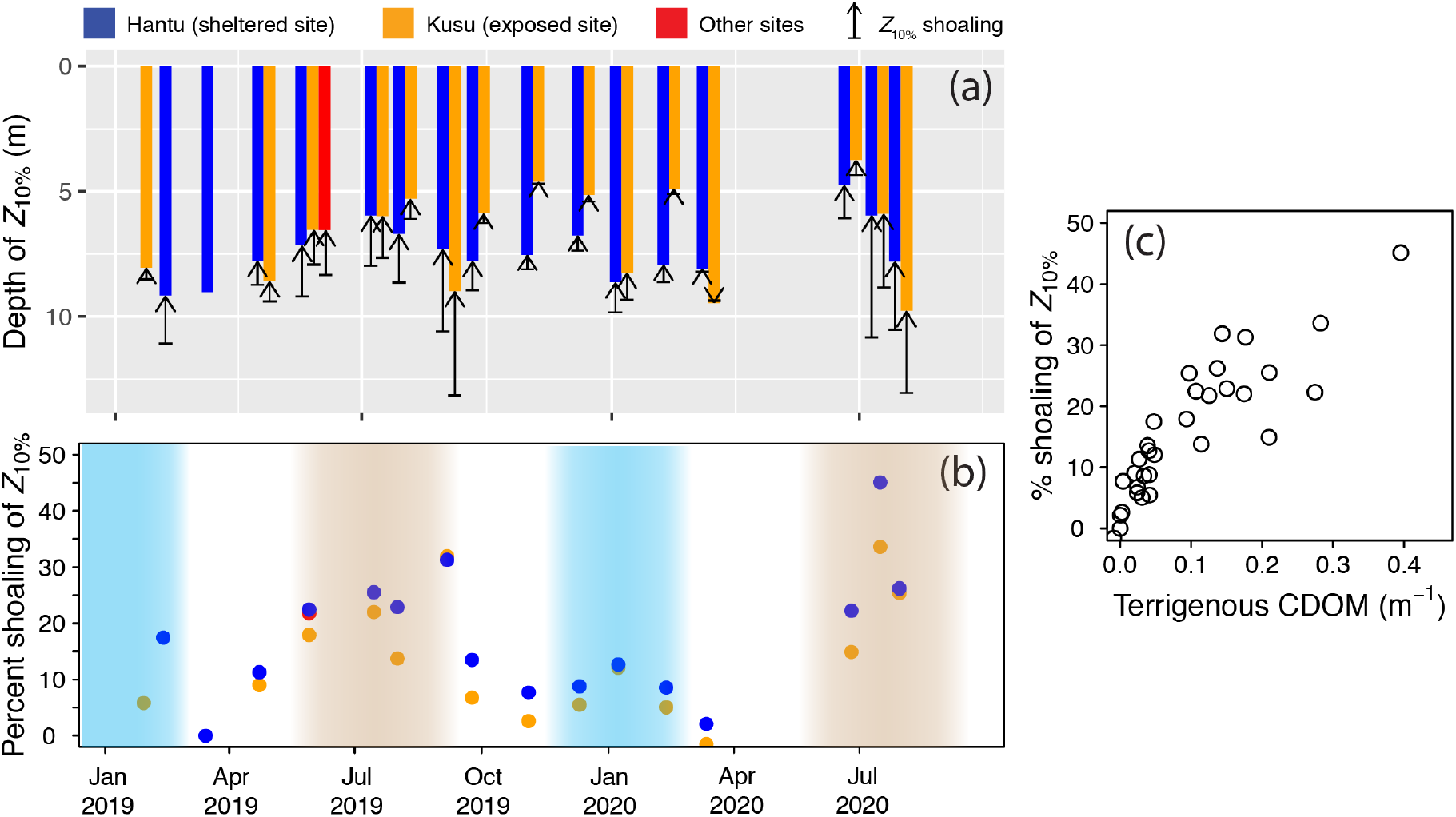
Time-series data showing the impact of terrigenous CDOM on the depth of 10% PAR penetration (*Z*_10%_). (a) Bars show the actually observed depth of *Z*_10%_, and arrows indicate by how much the depth of *Z*_10%_ was shoaled due to the presence of terrigenous CDOM (i.e., without terrigenous CDOM, *Z*_10%_ would extend to the bottom of the arrows). (b) Time series of the percentage reduction in *Z*_10%_ due to terrigenous CDOM. (c) Scatter plot showing the strong relationship between percentage reduction in *Z*_10%_ and the absorption coefficient at 440 nm by terrigenous CDOM.

To estimate how much of the observed shoaling of *Z*_10%_ might be the result of anthropogenic disturbance of peatlands, we repeated our calculations of *K*_*d*_ for the SW Monsoon but with the terrigenous CDOM absorption reduced by 35% of the observed value (see Section 2.8). We found that with only the estimated natural fraction of terrigenous CDOM, *Z*_10%_ was on average 0.6 m deeper than the actual observed values. This potentially anthropogenic contribution to light attenuation accounted for on average 25% of the observed *Z*_10%_ shoaling by terrigenous CDOM (Table 1).

**Table 1.** Estimated contribution to the shoaling of the depth of 10% PAR penetration (*Z*_10%_) by terrigenous CDOM released by land-conversion of peatlands. Because our sites only receive peatland-derived CDOM during the Southwest Monsoon, this estimate was only made for measurements collected during this season. The percentage anthropogenic shoaling was calculated relative to the amount of shoaling caused by the total observed terrigenous CDOM. The anthropogenic terrigenous CDOM fraction was estimated as 35% of observed terrigenous CDOM, based on previous work (Moore et al. 2013, Yupi et al. 2016).

| Date | Site | Latitude (°N) | Longitude (°E) | $Z_{10\%}$ shoaling due to anthropogenic terrigenous CDOM (m) | Anthropogenic contribution to total shoaling by terrigenous CDOM (%) |
| --- | --- | --- | --- | --- | --- |
| 2019-05-29 | Hantu | 1.227 | 103.746 | 0.56 | 27.2 |
| 2019-05-29 | Kusu | 1.226 | 103.860 | 0.40 | 28.1 |
| 2019-05-29 | Other | 1.246 | 103.738 | 0.48 | 26.4 |
| 2019-07-15 | Hantu | 1.227 | 103.746 | 0.50 | 24.4 |
| 2019-07-15 | Kusu | 1.226 | 103.860 | 0.45 | 26.3 |
| 2019-08-01 | Hantu | 1.227 | 103.746 | 0.52 | 26.3 |
| 2019-08-01 | Kusu | 1.226 | 103.860 | 0.24 | 27.9 |
| 2019-09-06 | Hantu | 1.227 | 103.746 | 0.79 | 23.7 |
| 2019-09-06 | Kusu | 1.226 | 103.860 | 1.00 | 23.8 |
| 2020-06-25 | Hantu | 1.227 | 103.746 | 0.33 | 24.1 |
| 2020-06-25 | Kusu | 1.226 | 103.860 | 0.18 | 27.1 |
| 2020-07-16 | Hantu | 1.227 | 103.746 | 0.89 | 18.2 |
| 2020-07-16 | Kusu | 1.226 | 103.860 | 0.65 | 21.9 |
| 2020-07-30 | Hantu | 1.227 | 103.746 | 0.71 | 25.6 |
| 2020-07-30 | Kusu | 1.226 | 103.860 | 0.87 | 26.1 |

## 4. Discussion

### 4.1 Sources and seasonality of CDOM

Our time series data showed strong and correlated seasonal variation in CDOM absorption, markers of CDOM terrestrial origin (S_275–295_, S_R_, and SUVA_254_), and salinity. Moreover, the variability in CDOM across all seasons followed a pattern that is consistent with simple conservative mixing between two CDOM end-members (Fig. 3), where one end-member has high *a*_CDOM_ and SUVA_254_ with low S_275–295_ and S_R_ (SW Monsoon) and the other has low *a*_CDOM_ and SUVA_254_ with high S_275-295_ and S_R_ (intermonsoon). The spectral slope S_275–295_ and the slope ratio S_R_ have become well established as accurate markers of terrigenous CDOM in regions receiving terrestrial inputs (Helms et al. 2008, Fichot & Benner 2012, Dainard & Guéguen 2013, Lu et al. 2016, Medeiros et al. 2017, Martin et al. 2018, Painter et al. 2018, Carr et al. 2019). Together with the correlated changes in SUVA_254_, our data therefore show that the Singapore Strait receives a large input of high-molecular-weight, aromatic-rich, terrigenous CDOM during the SW Monsoon, and a smaller input of terrigenous CDOM during the NE Monsoon. In contrast, the CDOM pool during the intermonsoon seasons appears to be characterised by low molecular weight and low aromaticity, consistent with a primarily marine, autochthonous source of CDOM such as from planktonic production. Importantly, our data show that the seasonal changes in CDOM absorption can be largely explained by conservative mixing of the intermonsoon marine CDOM with terrigenous CDOM delivered during the monsoon seasons. This justifies our interpretation that CDOM in the Singapore Strait is composed of a relatively constant and low background level of marine CDOM with a seasonally variable admixture of terrigenous CDOM.

Our conclusion that the CDOM during the SW Monsoon is predominantly terrigenous is further supported by the fact that the stable carbon isotope composition of DOC (*δ*^13^C_DOC_) reached values as low as −25.5‰ during this time, which can only be explained by a large, seasonal input of tDOC (Zhou et al. submitted manuscript). In contrast, the *δ*^13^C_DOC_ averaged around −22‰ during the late NE Monsoon and Intermonsoon 1 periods (Zhou et al. submitted manuscript), which is consistent with a total DOC pool that is fully of marine origin. The high salinities during the intermonsoon seasons (up to 33) are also close to typical values for the central South China Sea, which are mostly <34 (Wong et al. 2007), indicating that freshwater input (and therefore the potential for new input of terrigenous CDOM) is small outside of the two monsoon seasons. Because terrigenous CDOM in Southeast Asia also appears to be readily photo-bleachable (Martin et al. 2018), any terrigenous CDOM delivered to the Singapore Strait during the NE Monsoon is also likely to be removed by photo-bleaching during the intermonsoon period.

The fact that chlorophyll-*a* concentrations did not vary seasonally and were overall relatively low for a coastal environment further indicates that production of autochthonous, marine CDOM is unlikely to show strong seasonal variation (note that benthic communities such as coral reefs and seagrasses are restricted to small areas of the Singapore Strait (Tan et al. 2016), and are hence very unlikely to be quantitatively significant sources of autochthonous CDOM). The close similarity in the values of *a*_CDOM_(440) between the intermonsoon months in all three years (2018–2020, Fig. S2) provides additional evidence at interannual time-scales that the background level of marine CDOM does not vary strongly over time in the Singapore Strait.

Given both the seasonal pattern of ocean currents and the seasonally high values of CDOM absorption, peatlands on Sumatra are the only plausible source of the terrigenous CDOM during the SW Monsoon. This is consistent with satellite remote sensing data that show CDOM spreading out from Sumatra into the Malacca and Karimata Straits (Siegel et al. 2019). Although the smaller input of terrigenous CDOM during the NE Monsoon follows the same theoretical mixing line as the data from the SW Monsoon (Fig. 3), the differences in ocean currents obviously rule out Sumatran peatlands as the source of this CDOM. The terrigenous CDOM during the NE Monsoon might be derived largely from river input along the east coast of the Malay Peninsula (Kuwahara et al. 2010, Mizubayashi et al. 2013), which mostly consists of mineral soils rather than peatlands (Fig. 1).

### 4.2 Impact of terrigenous CDOM on light availability

The strong light attenuation, and resulting shallow *Z*_10%_ depth that we observed in our time series is consistent with previous reports of strong vertical attenuation of photosynthetically active radiation (PAR) in the Singapore Strait (Dikou & van Woesik 2006, Chow et al. 2019, Morgan et al. 2020). Our data additionally show that the seasonal input of terrigenous CDOM to the Singapore Strait clearly contributes significantly to the extinction of PAR with depth, and also alters the spectral quality of the available light. This was already evident when just considering the seasonal averages of *Z*_10%_, even though the observed *Z*_10%_ was also clearly affected by the variability in particulate absorption and backscattering. After first estimating the fraction of CDOM that was terrigenous, we could quantify its impact directly by calculating hypothetically how the spectrum of *K*_*d*_ would differ in its absence; this showed that the advection of terrigenous CDOM during both monsoon seasons leads to shoaling of *Z*_10%_ by tens of percent (Fig. 8). Moreover, this shoaling was accompanied by spectral shifts in underwater irradiance, leading to less blue light and an irradiance peak shifted towards longer wavelengths (Figs. 5,6).

This result is consistent with the known importance of CDOM in reducing light penetration and altering the spectral quality of light in coastal waters (DeGrandpre et al. 1996, Foden et al. 2008, Mascarenhas et al. 2017). In coral reefs specifically, CDOM exerts a major control over the attenuation of UV radiation (Dunne & Brown 1996, Otis et al. 2004, Zepp et al. 2008, Kuwahara et al. 2010), and reefs off Peninsular Malaysia have been shown to receive significant inputs of terrigenous CDOM (Kuwahara et al. 2010, Bowers et al. 2012).

Mizubayashi et al. (2013) further showed that this terrigenous CDOM input is correlated with changes in UV and PAR attenuation off north-eastern Malaysia. Our data thus provide further evidence of the importance of CDOM in controlling the light environment of coral reefs, but also demonstrate that the spectral distribution of PAR is affected.

Our results also provide further support for the use of CDOM spectral slope measurements, especially S_275–295_, to distinguish between marine and terrigenous CDOM in coastal waters (Stedmon & Markager 2001, Helms et al. 2008, Astoreca et al. 2009, Dainard & Guéguen 2013, Vantrepotte et al. 2015, Lu et al. 2016). Such a partitioning between marine and terrigenous CDOM fractions and their respective contributions to the spectral light attenuation in coastal waters is needed for a better understanding of the potential drivers of shelf sea light availability.

Our analysis also indicates that around 25% of the seasonal CDOM-mediated shoaling of *Z*_10%_ might be an anthropogenic effect caused by the increase in tDOC flux due to peatland disturbance (Moore et al. 2013, Yupi et al. 2016). Although it has been well documented that the CDOM pool in shelf seas can contain a large fraction of terrigenous CDOM (Blough et al. 1993, Stedmon et al. 2010, Mizubayashi et al. 2013, Carr et al. 2019), the long-term dynamics and potential anthropogenic drivers of terrigenous CDOM in coastal waters remain poorly known. So far, there is only limited and mostly indirect evidence that anthropogenic increases in coastal CDOM concentrations have occurred and reduced light penetration, and this has only been reported off southern Norway (Aksnes et al. 2009, Frigstad et al. 2013) and in the Gulf of Maine (Balch et al. 2016). Our estimate of the anthropogenic contribution to the observed CDOM-mediated light attenuation relies on earlier reports that the peatland tDOC flux has increased by slightly over 50% as a result of land conversion (Moore et al. 2013, Yupi et al. 2016). Future research should therefore aim to corroborate our estimate by reconstructing past variation in terrigenous CDOM in this region, which may become possible through measurements of humic acid concentrations in coral skeleton cores (Kaushal et al. 2020). Nevertheless, our study already indicates that the disturbance of tropical peatlands has likely resulted in CDOM-mediated coastal browning in Southeast Asia, and that peatland disturbance therefore entails an additional environmental impact beyond the large increases in CO_2_ emissions from peat and peatland DOC oxidation (Hooijer et al. 2010, Murdiyarso et al. 2010, Wit et al. 2018).

### 4.3 Ecological implications

Primary production by benthic communities and by phytoplankton requires sufficient light availability, and strong extinction of PAR can therefore limit productivity and restrict the depth to which photosynthetic benthic communities can occur (Gattuso et al. 2006). Moreover, because different phytoplankton taxa differ in their pigment composition and photo-acclimation strategies, the spectral quality of underwater irradiance can control phytoplankton community composition (Glover et al. 1987, Palenik 2001, Grébert et al. 2018). This has been shown specifically also for large, CDOM-driven spectral shifts from blue/green to red wavelengths (Stomp et al. 2004, Stomp et al. 2007, Frenette et al. 2012, Lawrenz & Richardson 2017).

Our estimates of the depth-averaged underwater irradiance in a turbulent water column show that phytoplankton in the Singapore Strait are subject to seasonal changes in intensity and spectral composition of irradiance (Fig. 6). The fact that the phytoplankton absorption spectral slope increased and the *a*_Phyto_(490):*a*_Phyto_(510) ratio decreased during the SW Monsoon suggests that the phytoplankton were adjusting their pigment composition in response to the changing light environment. Specifically, these data indicate a higher ratio of photoprotective to photosynthetic carotenoid pigments during the intermonsoon, and a lower ratio during the SW Monsoon (Eisner et al. 2003, Hickman et al. 2009). Given that the phytoplankton community in the Singapore Strait does not undergo major seasonal changes (Gin et al. 2000, Chénard et al. 2019), our data most likely reflect photo-acclimation by individual taxa rather than a taxonomic community shift.

Such chromatic adaptation of marine phytoplankton is known from coastal and open-ocean environments, as a function of both vertical and horizontal variation in the light environment (Hickman et al. 2009, Isada et al. 2013, Pérez et al. 2020). Whether the changes we observed in the *a*_Phyto_ spectra were driven more by the seasonal change in spectral light quality or more by the overall reduction in underwater PAR is unclear, but they indicate that the seasonal change in light availability caused by peatland CDOM was sufficiently large to require changes in photo-acclimation by the phytoplankton community. Lower light availability during the SW Monsoon may be a reason for why the chlorophyll-*a* concentration remains relatively constant (Fig. S4) despite increases in nutrient concentrations during this season by 3–5 μmol l^−1^ DIN and 0.3–0.4 μmol l^−1^ DIP (Chénard et al. 2019).

Our data also show that benthic communities within the upper 10 m are exposed to seasonal changes both in total PAR intensity and in spectral quality of irradiance (Figs. 5,8). The Singapore Strait is home to >100 different scleractinian coral species, but their depth range is restricted to within the upper 10 m (Huang et al. 2009). This shallow depth distribution is attributed chiefly to a combination of sediment stress and light limitation (Dikou & van Woesik 2006, Guest et al. 2016, Chow et al. 2019, Morgan et al. 2020), and matches quite closely with the average *Z*_10%_ of around 7 m that we measured. Sedimentation and light limitation together appear to have driven a process of vertical reef compression (Morgan et al. 2020), with coral reef monitoring data suggesting that coral cover at depths of 6–7 m (but not at 3–4 m) has decreased since the 1980s (Guest et al. 2016).

It has been suggested that coral communities in shallow low-light environments such as Singapore should be considered as mesophotic coral ecosystems, like those found below 30–40 m depth in clear, “blue-water” environments (Laverick et al. 2020, Morgan et al. 2020). Although these systems do show many ecological similarities, e.g., in species composition (Eyal et al. 2016, Chow et al. 2019, Laverick et al. 2020), the spectral distribution of the available light in Singapore is clearly very different compared to deep mesophotic ecosystems that receive mostly blue wavelengths (Kahng et al. 2019). Whether such differences in spectral light quality, and the additional seasonal changes we report here, are physiologically and ecologically significant for coral reefs is unclear. Corals can clearly acclimate to low light intensities both in deep and shallow mesophotic conditions (see review by Kahng et al. (2019)), such as by altering their skeletal morphology to minimise self-shading (Todd 2008, Ow & Todd 2010), and by enhancing light scattering by the skeleton and light absorption within the tissue layer (Polinski & Voss 2018, Kramer et al. 2020). However, the spectral light quality does appear to affect coral physiology. For example, calcification rates of two coral species were substantially increased under blue light compared to other wavelengths for equal light intensities (Cohen et al. 2016), and blue light acclimation increased maximum photosynthetic rates in *Montipora verrucosa* relative to green and red light (Kinzie & Hunter 1987). Branches of *Stylophora pistillata* acclimated to either blue or full-spectrum light showed better photosynthetic performance under the spectral conditions they were acclimated to (Mass et al. 2010). Blue light also controls the fluorescent pigmentation of several coral taxa (D’Angelo et al. 2008), while red light was found to reduce the health and survival of *Stylophora pistillata* (Wijgerde et al. 2014). Whether fluorescent proteins play a photo-physiological role in enabling coral acclimation to low-intensity blue light is still debated (D’Angelo et al. 2008, Roth et al. 2015, Smith et al. 2017, Kahng et al. 2019). However, the presence of photoconvertible red fluorescent proteins was necessary for long-term survival of two coral species under low-intensity blue light (Smith et al. 2017). It is therefore possible that the spectral differences in irradiance between shallow and deep mesophotic systems will prove to be ecologically significant, perhaps by controlling lower depth limits and coral community composition, or requiring a greater reliance on heterotrophic *versus* autotrophic nutrition (Anthony & Fabricius 2000).

The decline in coral cover at 6–7 m in the Singapore Strait since the 1980s reported by Guest et al. (2016) was originally attributed to possible increases in suspended sediments and sedimentation. However, this period of coral cover loss at deeper sites also coincides with the major period of land conversion of peatlands across Southeast Asia (Miettinen et al. 2016). Our results indicate that if peatland disturbance has indeed increased tDOC fluxes by as much as currently thought, then the associated reduction in light transmission due to terrigenous CDOM has likely contributed to these benthic cover changes.

## 5. Conclusions

Our data demonstrate the importance of terrigenous CDOM for the optical properties of peatland-influenced areas of the Sunda Shelf Sea, and show further that the seasonal, monsoon-driven advection of this terrigenous CDOM drives significant variation in the transmission and spectral quality of light underwater. We also observed seasonal variation in phytoplankton absorption spectra that are indicative of changes in photo-acclimation, which suggests that this variation in light attenuation was ecologically relevant. Moreover, our study suggests that land conversion in the tropics has the potential to cause CDOM-mediated coastal browning in biodiverse shelf sea environments, which may have contributed to observed coral cover decline. Overall, our study underscores the importance of examining not only biogeochemical impacts of land–ocean tDOC fluxes, but also the consequences for optical water quality due to the associated terrigenous CDOM.

## Acknowledgements

Yongli Zhou, Chen Shuang, Nikita Kaushal, Molly Moynihan, Rob Nichols, Tan Li, Daniel Kalbermatter, Jervis Ong Zhe Ao, Lee Tian Li, Phyllis Kho Yu Yi, Kyle Morgan, Woo Oon Yee, and Chen Yuan assisted with fieldwork and laboratory analyses. We thank Sapari, Surpato, and Francis Yeo of *Dolphin Explorer* for enabling the sample collection. Comments by Richard Sanders, Adam Switzer, and three anonymous reviewers improved this manuscript. Field work was carried out under permit NP/RP17-044-2 from the Singapore National Parks Board. This research was supported by the National Research Foundation Singapore, Prime Minister’s Office, under the Marine Science Research and Development Program through grant MSRDP-P32 to P.M.

## Author contributions

Conceptualization: P.M., N.S. and E.W.W.; Investigation and Formal analysis: P.M., N.S, T.W.Q.L, K.C, J.M.C.W and E.W.W; Writing – Original draft: P.M.; Writing – Review and editing: all authors; Supervision: P.M. and S.C.L.; Funding acquisition and Project administration: P.M.

## Data availability

The final dataset used in this study, all data analysis codes, and all raw data are available *via* the NTU Data Repository under doi: 10.21979/N9/TXYRC3.

