## Supplementary text and figures for "Dissolved organic matter from tropical peatlands impacts shelf sea light availability on coral reefs in the Singapore Strait, Southeast Asia"

**Validation of $K_d$ and $Z_{10\%}$ calculation using radiometer measurements**

At 19 of the 32 stations at which all absorption and backscattering parameters were measured, depth profiles of downwelling irradiance ($E_d$) were collected with a TriOS RAMSES ACC-Vis radiometer to measure the vertical attenuation coefficient of downwelling irradiance, $K_d$, and to estimate the depth at which downwelling PAR was reduced to 10% of surface PAR($Z_{10\%}$). These measurements were collected in order to verify that the calculated light attenuation spectra from absorption and backscatter measurements were accurate. The radiometer data also allowed us to establish a simple empirical relationship between $Z_{10\%}$ and $K_d$ at 520 nm (Fig. S1), which we used to determine the depth to which to calculate $K_d$(PAR) from the $K_d$ spectra, as explained in Section 2.6 of the main text.

The radiometer was deployed on the sun side of the boat, and pushed approximately 1.5 m away from the boat using a pole with a pulley, and lowered to between 6.0–8.0 m over the course of 2–3 minutes. The reference sensor was mounted upright above the top of the boat and away from any obstructions such as communication aerials. Data were acquired every 4 s at 2-nm resolution from 318–950 nm, and processed into final, calibrated spectra using the TriOS software. Data were then exported and further analyzed in R to calculate $K_d$ as the slope of a linear regression of the natural log of downwelling irradiance ($E_d$) *versus* depth. In some cases, surface focusing of sunlight caused high variability in measured $E_d$ within the upper 1–1.5 m. In such cases, the surface data were removed, although this did not strongly affect the resulting $K_d$ estimates. To assess the quality of the $K_d$ estimate, we examined the $r^2$ values of the linear regression models at every wavelength, which were always higher than 0.84, and mostly higher than 0.90. The spectrum of $E_d$ for each individual radiometer measurement was also converted to a total photon flux between 400–700 nm and $K_d$(PAR) calculated as the slope of a linear regression of the natural log of downwelling PAR *versus* depth. The depth of 10% PAR penetration was then calculated as 2.303/$K_d$(PAR).

The $K_d$ spectra measured with the radiometer agreed closely with the $K_d$ spectra calculated from absorption and backscattering, except for two outliers, with a root mean squared error <0.06 $m^{-1}$ at most wavelengths and mean absolute percent error <12% (Fig. S5). The measured and calculated value of $Z_{10\%}$ also agreed well, with root mean square error of 0.84 m and mean absolute percent error of 10% (Fig. S6). For two stations, the measured $K_d$ spectra differed from the calculated $K_d$ spectra by 0.2–0.4 $m^{-1}$ (grey points in Fig. S5a). It is possible that these two radiometer $K_d$ measurements were problematic, e.g. owing to accidental boat shadow effects, and these two stations were therefore omitted from the comparison.

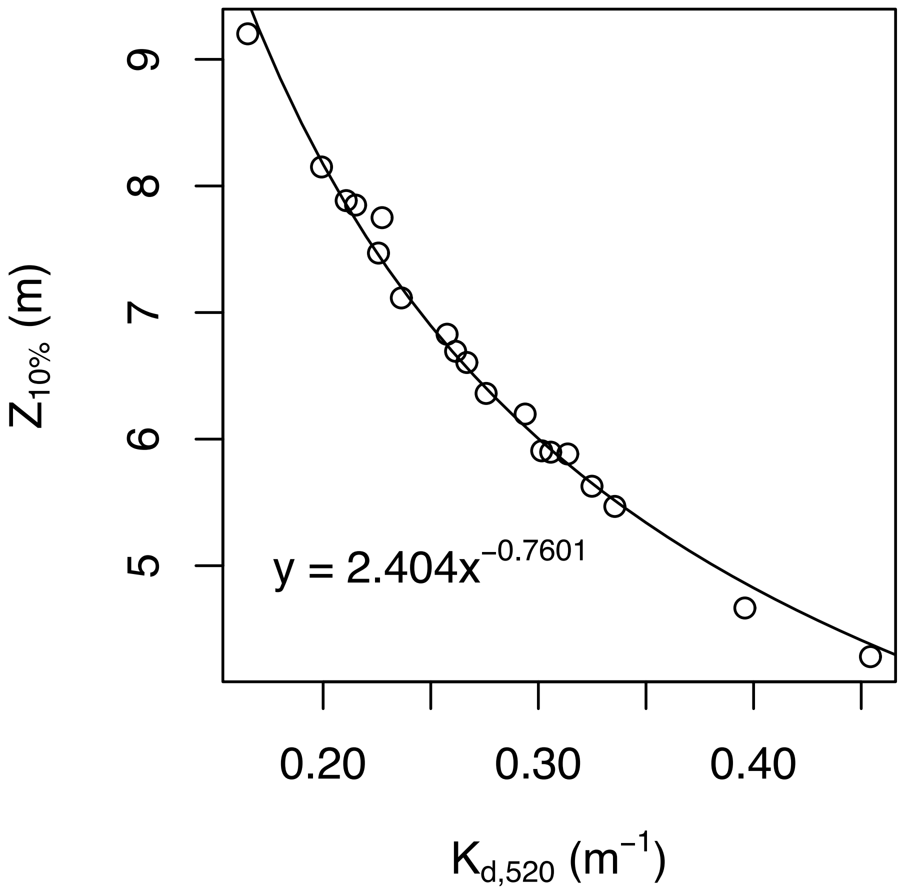


Fig. S1. The measured depth of 10% PAR penetration ($Z_{10\%}$) was closely related to $K_d$ at 520 nm with a power-law function (both measured with RAMSES radiometer). This relationship was used to determine the correct integration depth to calculate $K_d$(PAR) from the calculated $K_d$ spectra throughout our time series.

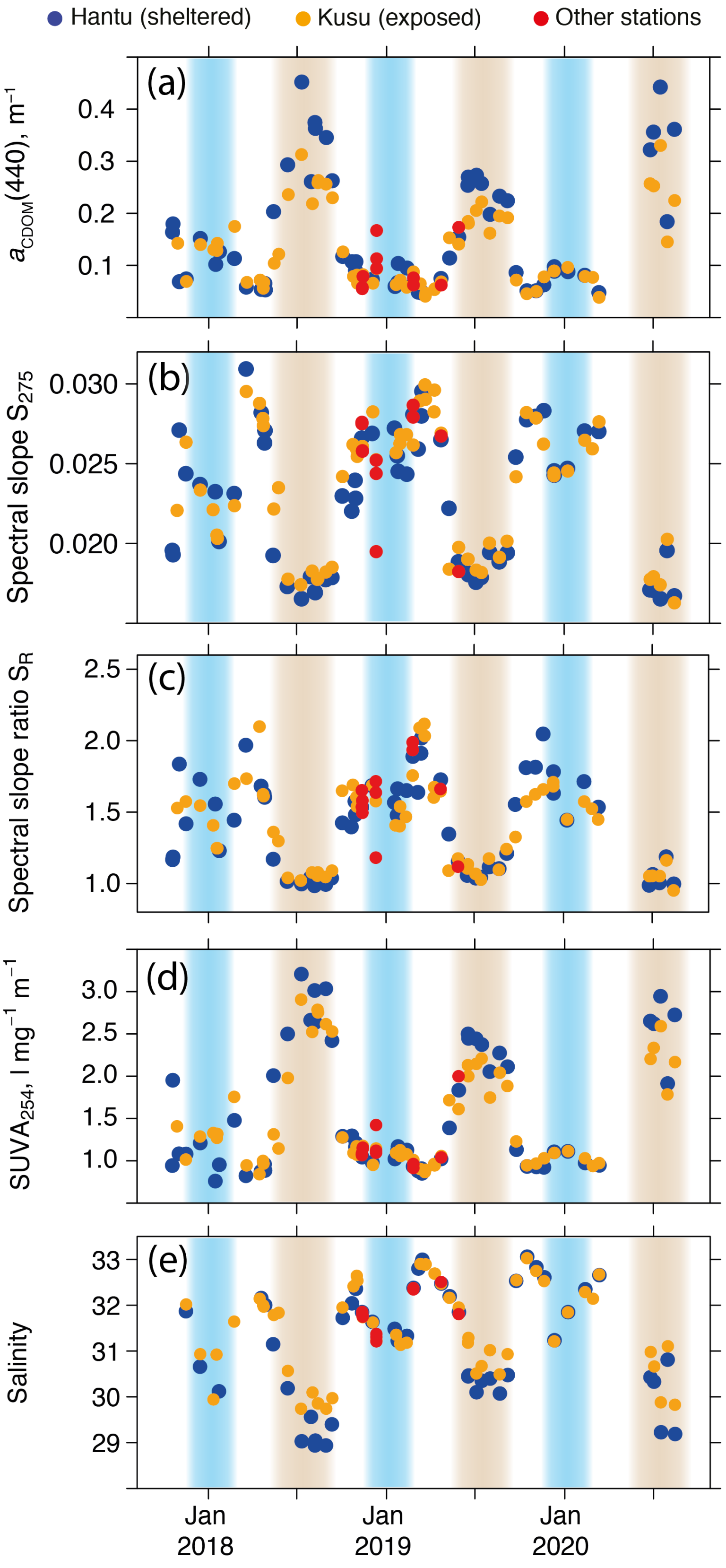


Fig. S2. Full time series from October 2017 to early August 2020 of (a) CDOM concentration (as CDOM absorption coefficient at 440 nm), (b) CDOM spectral slope from 275–295 nm, (c) the CDOM spectral slope ratio $S_R$, (d) the specific UV absorbance $SUVA_{254}$, and (e) seawater salinity. The SW and NE Monsoon seasons are marked with brown and blue shading, respectively.

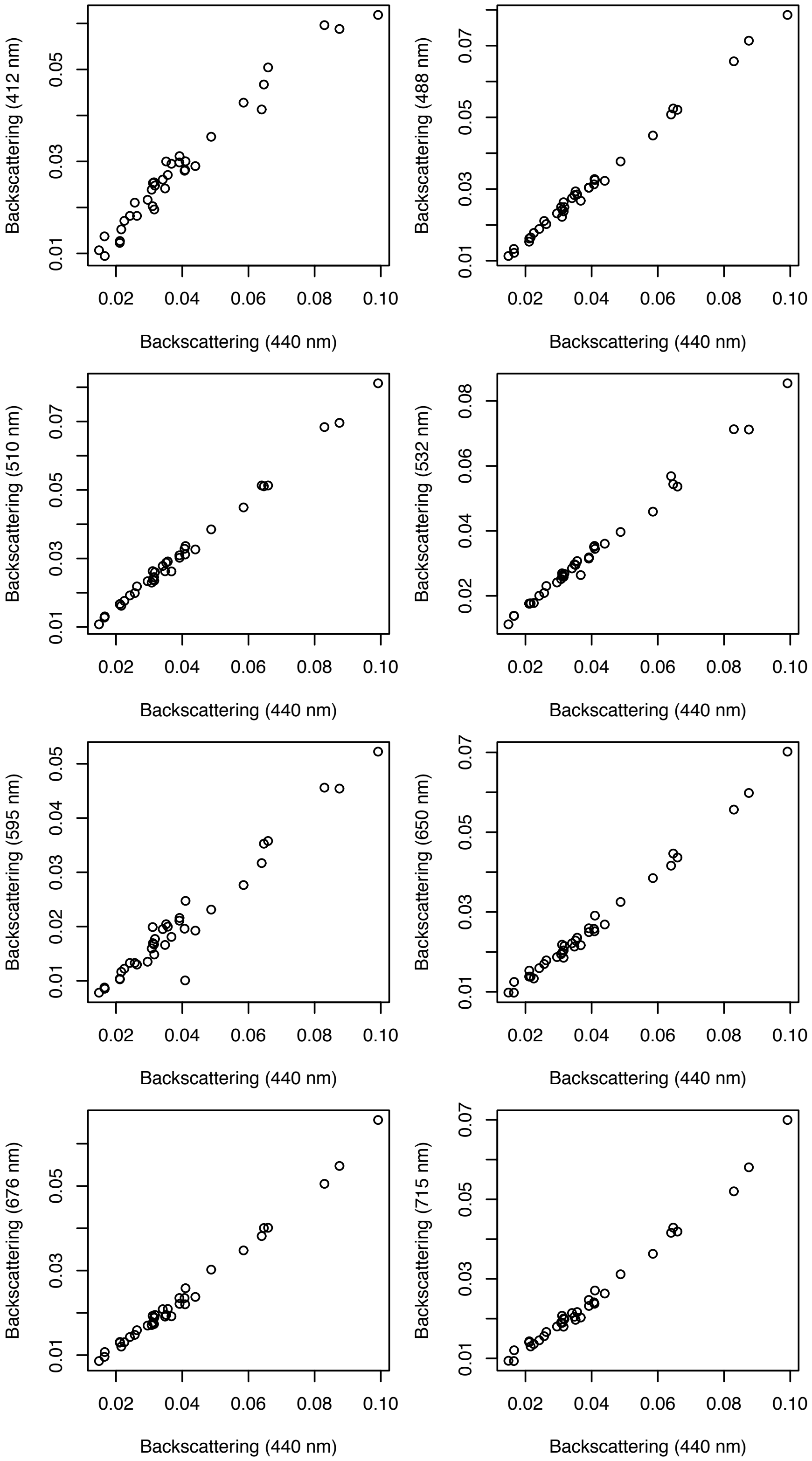


Fig. S3. Scatter plots showing that particulate backscattering at all other eight wavelengths was very closely related to particulate backscattering at 440 nm.

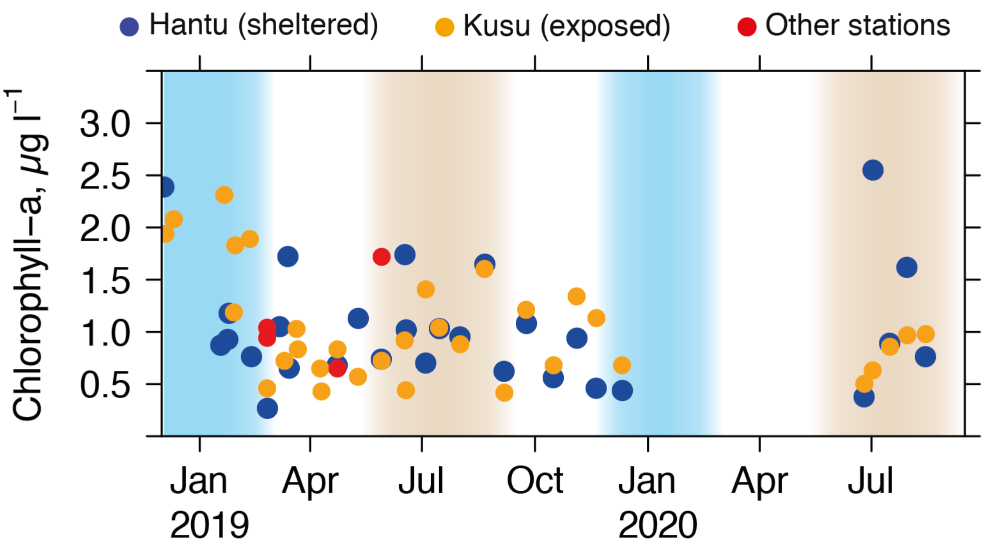


Fig. S4. Time series of chlorophyll-*a* concentration. The SW and NE Monsoon seasons are marked with brown and blue shading, respectively.

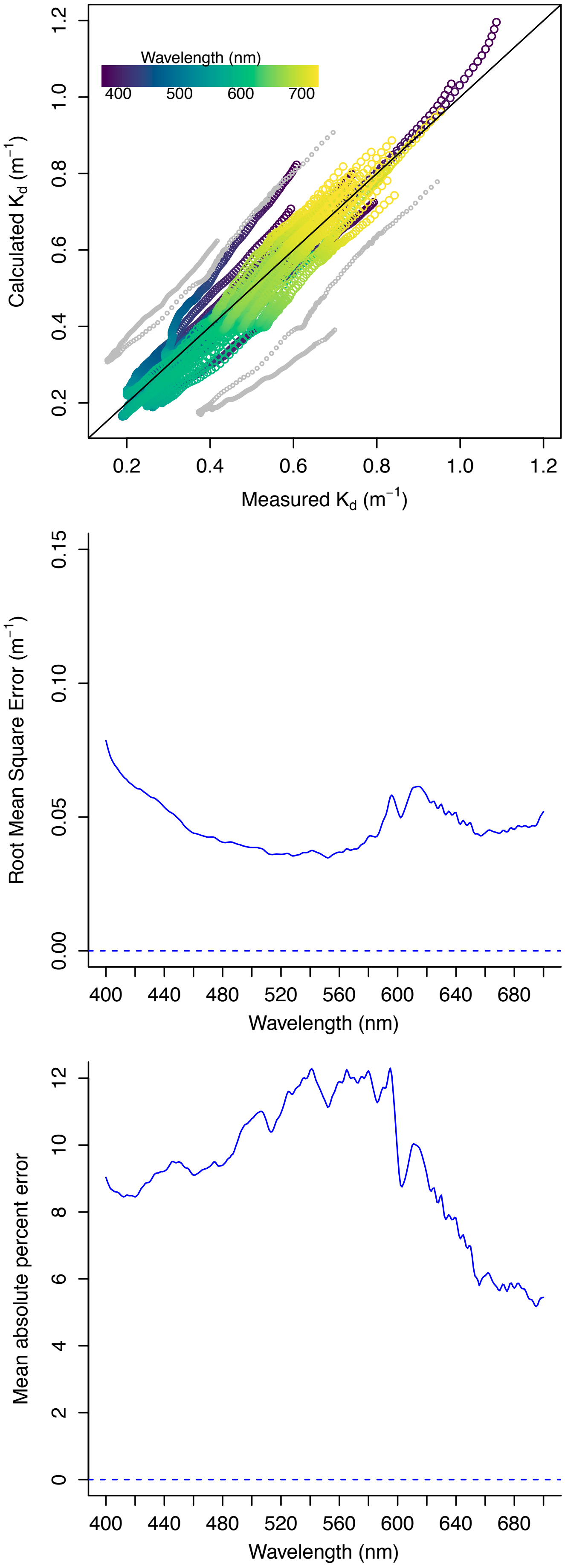


Fig. S5. (a) Calculated $K_d$ according to Lee et al. (2005) *versus* measured $K_d$ from TriOS RAMSES radiometer (n = 19 spectra). For each station, the full spectrum of $K_d$ is shown, with point colors indicating wavelength. Black line shows 1:1 relationship. The two outlier spectra shown in grey were omitted from the comparison between measured and calculated $K_d$. (b) Root mean square error between calculated and measured $K_d$ at all wavelengths (n = 17). (c) Mean absolute percent error between calculated and measured $K_d$ at all wavelengths (n = 17).

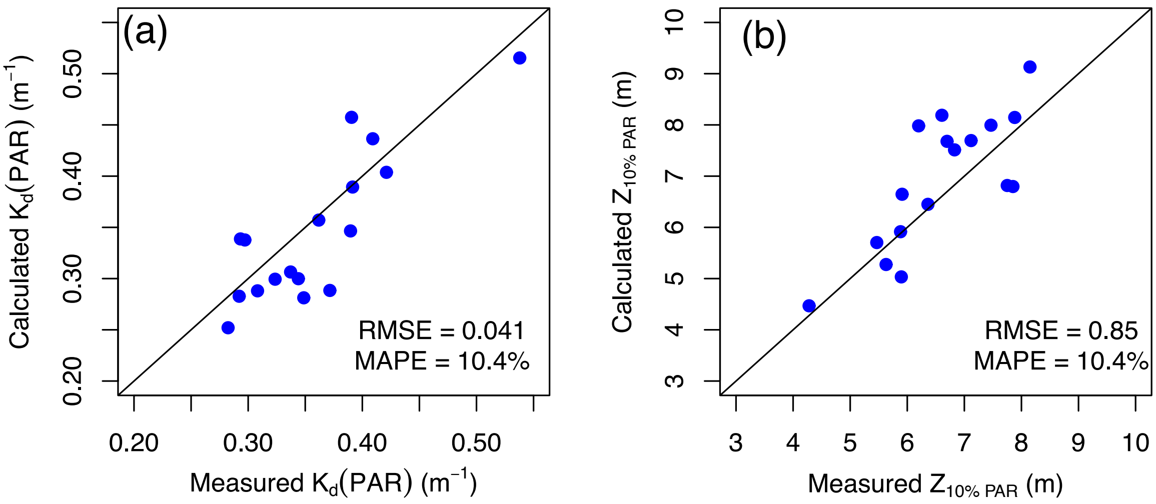


Fig. S6. Calculated (a) $K_d$(PAR) and (b) depth of 10% PAR penetration *versus* measurements taken with TriOS RAMSES radiometer (n = 17). Black lines show the 1:1 relationship. RMSE = root mean square error; MAPE = mean absolute percent error. Two outliers were removed from this comparison, as for Fig. S5.
